# Age-related structural and functional changes of the intracardiac nervous system

**DOI:** 10.1101/2023.11.24.568538

**Authors:** Eliza Sassu, Gavin Tumlinson, Dragana Stefanovska, Marbely C. Fernández, Pia Iaconianni, Josef Madl, Tomás Brennan, Manuel Koch, Breanne A. Cameron, Sebastian Preissl, Ursula Ravens, Franziska Schneider-Warme, Peter Kohl, Callum M. Zgierski-Johnston, Luis Hortells

## Abstract

**Background:** Although aging is known to be associated with an increased incidence of both atrial and ventricular arrhythmias, there is limited knowledge about how Schwann cells (SC) and the intracardiac nervous system (iCNS) remodel with age. Here we investigate the differences in cardiac SC, parasympathetic nerve fibers, and muscarinic acetylcholine receptor M2 (M2R) expression in young and old mice. Additionally, we examine age-related changes in cardiac responses to sympathomimetic and parasympathomimetic drugs.

**Methods and Results:** Lower SC density, lower SC proliferation and fewer parasympathetic nerve fibers were observed in cardiac and, as a control sciatic nerves from old (20–24 months) compared to young mice (2–3 months). In old mice, CSPG4 was increased in sciatic but not cardiac nerves. Expression of M2R was lower in ventricular myocardium and ventricular conduction system from old mice compared to young mice, while no significant difference was seen in M2R expression in sino-atrial or atrio-ventricular node pacemaker tissue. Heart rate was slower and PQ intervals were longer in Langendorff-perfused hearts from old mice. Ventricular tachycardia and fibrillation were more frequently observed in response to carbachol administration in hearts from old mice versus those from young mice.

**Conclusions:** On the background of reduced presence of SC and parasympathetic nerve fibers, and of lower M2R expression in ventricular cardiomyocytes and conduction system of aged hearts, the propensity of ventricular arrhythmogenesis upon parasympathomimetic drug application is increased. Whether this is caused by an increase in heterogeneity of iCNS structure and function remains to be elucidated.

**Graphical abstract:** 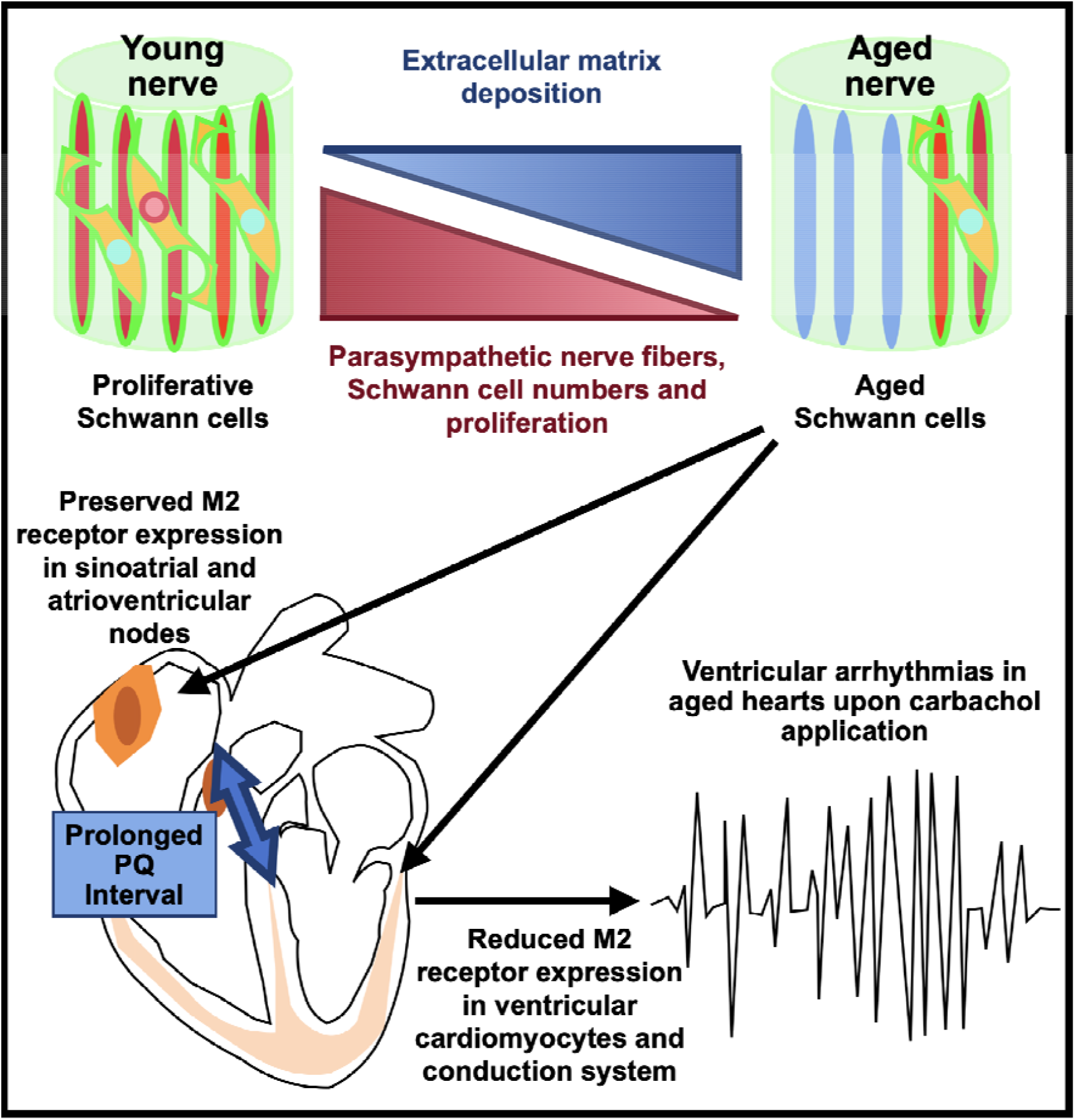

## 1. Introduction

Aging is characterized by structural tissue remodeling that is often associated with altered organ function and increased vulnerability to disease [1]. While aging-associated remodeling of the peripheral nervous system (PNS) has been studied in the context of skeletal muscle dysfunction [2], the relationship between age-related remodeling of the intracardiac nervous system (iCNS) and cardiac arrhythmogenesis – a marked problem associated with aging – is insufficiently understood.

PNS and iCNS are composed of sympathetic, parasympathetic, sensory, and intercircuit neurons, which are supported by myelinating Schwann cells (SC), non-myelinating Remak SC, endoneurial fibroblasts, microglia, macrophages, and vascular cells [3–5]. Whereas the distribution and function of the different cell populations in the PNS are well described, their iCNS counterparts have not been thoroughly characterized. In particular, few studies have focused on cardiac SC: one study demonstrated their presence in the heart using electron microscopy [5], while another described their role in postnatal cardiac maturation [3]. Importantly, electrophysiological abnormalities can be induced by decreasing the number of postnatal SC, which points to their potential relevance in iCNS-maturation and heart rhythm maintenance [6].

One of the main functions of PNS SC is to provide metabolic support to neuronal axons. In the PNS, age-related changes in the expression of proteins involved in SC proliferation and death have been associated with a reduced ability to repair nerves [7]. Degeneration and retraction of nerve fibers during aging has been described for the PNS [8] and the iCNS [9]. In the heart, aging has been associated with sympathetic neurodegeneration, which may be associated with increased β-adrenoreceptor sensitivity, while parasympathetic remodeling of the iCNS during aging remains underexplored.

Imbalanced iCNS function has been suggested to play a role in the development of atrial fibrillation [10] and myocardial infarction-related arrhythmias [11]. Recent findings point to a correlation between the release of the SC-derived S100 calcium-binding protein B (S100B) and cardiac neuronal damage in atrial ablation patients [12], but, differences in the numbers of S100B-expressing (S100B+) SC from old and young hearts have not been reported.

Here we describe changes in the PNS and iCNS that are associated with aging. In particular, we reveal age-related differences in cardiac SC presence and proliferation, report a reduction in cardiac parasympathetic nerve density, and illustrate coinciding changes in heart rate and rhythm.

## 2. Methods

### 2.1 Organ extraction from mice

All mouse experiments were carried out according to the guidelines stated in Directive 2010/63/EU of the European Parliament on the protection of animals used for scientific purposes and were approved by the local authorities (Regierungspräsidium Freiburg; G16/131) and by the University of Freiburg (X19-01R).

Experiments were performed on tissue obtained from young (age: 2–3 months) or old (age: 20–24 months) C57BL/6J (WT) mice, unless stated otherwise. *Tcf21*-MerCreMer (*Tcf21^MCM^*) [13] and alpha myosin heavy chain Cre (*MHC^Cre^*; B6.FVB-Tg(Myh6-cre)2182 Md/J; Jax reference: 009074) mice were bred with channelrhodopsin 2-YFP mice (*R26^ChR2-YFP^*; 129S-Gt(ROSA)26Sortm32(CAG-COP4∗H134R/EYFP)Hze/J; Jax reference: 024109) mice to genetically trace cardiac fibroblasts or cardiomyocytes (CM), respectively. To induce the activity of the MerCreMer protein, intraperitoneal tamoxifen injections (3 mg/kg body weight on five consecutive days) were used, and hearts were harvested 14 days after the initial injection. Mice were euthanized by cervical dislocation, their chests were opened, and their hearts swiftly removed and placed in warm (37 °C) followed by cold (4 °C) heparin-containing (10 units/mL) ‘Tyrode solution’, containing [in mM]: 140 NaCl, 6 KCl, 10 HEPES, 10 glucose, 1.8 CaCl_2_, 1 MgCl_2_, pH titrated with 1 M NaOH to 7.4, 300±5 mOsmol/L. Hearts were either immediately used for functional studies or further processed for structural investigations by immunofluorescence imaging. Femoral sciatic nerves were sectioned from the sciatic notch to the trifurcation, by medial access through the biceps femoris in the posterior extremities. Cardiac nerves were identified [14, 15] and isolated from excised preparations of whole hearts (Fig S1). Nerve branches entering the cardiac parenchyma were visualized using magnifying lenses and were freed from surrounding tissue. The sino-atrial node (SAN) was dissected as previously described [16]. Nerves and SAN were washed and kept in phosphate-buffered saline (PBS) at 4 °C.

### 2.2 Immunohistochemical analyses and TUNEL assay

#### Whole-mount staining of the SAN and cardiac and sciatic nerves

Freshly harvested SAN, as well as cardiac and sciatic nerves, were fixed with 4% paraformaldehyde in PBS at 4 °C for 1 h. Thereafter, the tissue was washed three times in PBS and stored at 4 °C. Nerves were cut into three sections before staining; all staining steps were conducted at room temperature (RT). Unspecific antibody binding-site blocking and cell permeabilization were performed simultaneously by incubation for 1h in PBS containing 6% donkey serum and 0.25% Triton X-100. Incubation with primary antibodies was performed for 48 hours. After three washing steps, samples were incubated with Alexa-Fluor-conjugated (488, 555, 594, 647) donkey secondary antibodies (Abcam, Cambridge, UK) directed at the species in which primary antibodies were raised (see Table 1) at 1:400 dilution 48 hours. Nuclei were stained overnight with 4′,6-diamidino-2-phenylindole (DAPI) at 1:500 dilution in PBS. Finally, samples were mounted between two coverslips by embedding the tissue in Invitrogen ProLong Glass Antifade mounting medium (Thermo Fisher Scientific, Waltham, MA, USA) to match the refractive index of glass before being cured overnight at RT in a humidity-controlled environment.

**Table 1:** Primary antibodies utilized in this study.

| **Target Antigen** | **Host species** | **Provider** |
| --- | --- | --- |
| Green Fluorescent Protein (GFP)<br>1:100 | Goat | ab6673<br>Abcam, Cambridge, UK |
| Acetylcholinesterase (ChAT)<br>1:100 | Rabbit | ab183591<br>Abcam, Cambridge, UK |
| Tyrosine Hydroxylase (TH)<br>1:100 | Rabbit | ab112<br>Abcam, Cambridge, UK |
| SRY-Box Transcription Factor 10 (SOX10)<br>1:100 | Rabbit | ab180862<br>Abcam, Cambridge, UK |
| S100 calcium-binding protein B (S100B)<br>1:100 | Rabbit | ab52642<br>Abcam, Cambridge, UK |
| Histone3 (phosphor S10) (PH3)<br>1:100 | Mouse | ab14955<br>Abcam, Cambridge, UK |
| Muscarinic Acetylcholine Receptor 2 (M2R)<br>1:100 | Rabbit | ab109226<br>Abcam, Cambridge, UK |
| Hyperpolarization-activated cyclic nucleotide-gated channel 4 (HCN4)<br>1:100 | Rat | ab32675<br>Abcam, Cambridge, UK |
| Cluster of Differentiation 31 (CD31)<br>1:50 | Rat | 550274<br>BD biosciences, Haryana, India |
| Cluster of Differentiation 68 (CD68)<br>1:100 | Mouse | 153311<br>BIORAD, Hercules, CA, USA. |
| Chondroitin sulfate proteoglycan 4 (CSPG4)<br>1:200 | Rabbit | ab129051<br>Abcam, Cambridge, UK |
| Beta 3 tubulin (TUBB3)<br>1:500 | Mouse | ab78078<br>Abcam, Cambridge, UK |

#### Heart sections

Freshly harvested hearts were cut longitudinally into two pieces (exposing a four-chamber view, cut from the outflow tract to the apex), washed in PBS, embedded next to each other with the dorsal half epicardium facing down and ventral half endocardium facing down, in Sakura Finetek Tissue-Tek optimal cutting temperature compound (OCT; Thermo Fisher Scientific), and plunge-frozen in liquid nitrogen. 10 μm sections were cut for immunofluorescence imaging (at least four sections per heart were used for quantification). Slices were fixed for 15 min in 2% paraformaldehyde (Thermo Fisher Scientific) and permeabilized with 0.1%Triton X-100 (Thermo Fisher Scientific) for 15 min, all at RT. Unspecific antibody binding was blocked with 6% donkey serum or 4% goat serum (Sigma-Aldrich, St Louis, MO, USA), and 2.5% bovine serum albumin (BSA; MilliporeSigma, Burlington, MA, USA) for 1 h at RT. Samples were incubated overnight with primary antibodies at 4°C. After washing, samples were incubated for 1h with Alexa-conjugated (488, 555, 594, and 647) donkey or goat secondary antibodies at 1:400 dilution at RT (see Table 1). DAPI was applied at 1:500 dilution in PBS for 10 min at RT. Samples were mounted with Fluoromount-G mounting medium (Thermo Fisher Scientific).

#### TUNEL assay

To detect cell death, the terminal deoxynucleotidyl transferase dUTP nick end labelling (TUNEL) *in situ* cell death detection kit (Roche, Basel, Switzerland) was used according to the manufacturer’s instructions (including incubation for 1 h at 37 °C). When TUNEL and immunofluorescence were combined, the TUNEL reaction was performed after incubating the samples with primary antibodies and the secondary antibodies were added to the kit reaction buffer.

#### Microscopy

Fluorescence imaging was performed using a confocal microscope (TCS SP8 X, Leica Microsystems, Wetzlar, Germany) or a Fluorescence Microscope (Thunder Imager Leica Microsystems). All the images had an 8 bits size. For whole-mount immuno-stained sciatic nerves, six z-stack planes (2 - 6 µm step distance, 1024x1024 8-bit pixels) planes of the whole-nerve section were imaged, and nerve diameters were measured at their widest point. For mouse hearts, at least six longitudinal four-chamber mid-stack sections per heart were imaged and an overview cross-section of the left and right atria, as well as of the cardiac ventricles (left and right ventricular walls and septum), were imaged separately. At least eleven images (at 40x magnification) from each heart were analyzed for quantification of muscarinic acetylcholine receptor M2 (M2R) expression in SAN and ventricles.

### 2.3 Histological preparation and imaging of the sciatic nerve

Sciatic nerves were harvested, fixed and washed as described above, dehydrated in rising concentrations of ethanol (50%, 70%, 80%, 95%, 100%, 100%, and 100%, with 1 h incubation at each concentration), and incubated for 3 x 1 h in Xylene in an automatized system (TP1020 Tissue Processor, Leica Biosystems). Sciatic nerves were then embedded in paraffin and 5 µm sections were obtained with a microtome (Leica Biosystems). Paraffin sections were melted at 60 °C for 30 min, then deparaffinized and re-hydrated. Afterwards, a VitroView Alcian Blue Hematoxylin/Orange G Stain Kit (VitroVivo Biotech, Rockville, MD, USA) was used according to the manufacturer’s protocol. Samples were imaged with an automated slide scanner (AxioScan.Z1, Zeiss, Jena, Germany).

### 2.4 Image analysis

SOX10+/DAPI+ nuclei and S100B+ /DAPI+ nuclei cells were assumed to represent SC and counted using the Pixel Classification + Object Classification workflow in ilastik [17], an open-source, interactive machine learning platform for (bio)image analysis. Prebuilt Python packages (LifFile, numpy, and h5py) were used to extract data contained in proprietary formats. Five images of the sample types ‘heart’ or ‘sciatic nerve’ were chosen as a training set, lumping together tissue from young and old mice to consistently establish the algorithm. Pixel features, i.e., calculated values used to train ilastik to classify and group pixels into objects, were selected according to their presumed relevance in detecting cell nuclei. Preference was given to fluorescence intensity and edge over texture features as nuclei appeared as bright, sharply delineated, homogeneous objects. As each individual z-plane provided information on cell-type density, ilastik was set to forego computationally expensive 3D pixel grouping across z-planes. Manually highlighted nuclei (DAPI+ signal) were used to train ilastik to return all nuclei as individual objects, clearly separated from the background’. Ilastik was trained to recognize S100B+ or SOX10+ cells in multiple images, based on manual selections. Once ilastik could accurately and reliably highlight S100B+ or SOX10+ cells (assumed to comprise all SC) in a set of new, untrained images, the remaining dataset was automatically processed. Cell identities were exported and the ratio of SC to DAPI+ nuclei calculated.

M2R density was estimated as the proportion of cell membrane positive pixels that were also positive for M2R. Thresholds to eliminate background fluorescence were manually set and, along with imaging parameters, maintained for all data to allow comparison. A zero-filled array was generated for an absolute map of positive pixels. CSPG4 density in the sciatic nerve was estimated as the proportion of positive pixels against a zero-filled array (absolute map of positive pixels). In the heart, CSPG4 density was estimated as the proportion of the pan-neuronal marker Beta 3 Tubulin (TUBB3) [3] positive pixels that were also positive for CSPG4. Thresholds to eliminate background fluorescence were manually set and, along with imaging parameters, maintained for all data to allow comparison.

### 2.5 Whole-heart electrophysiological experiments

Hearts were excised as described above, weighed and the aorta cannulated (mean 357 s for young mice and 438 s for old mice) before transfer to a portable Langendorff perfusion system for transport to the measurement equipment. Here, hearts were perfused with oxygenated Tyrode solution at a flow rate of 3.3 mL per minute. Hearts were loaded with 15 μL of 1.4 mM Di-4-ANBDQPQ (dissolved in ethanol; Cytovolt2, CytoCybernetics, Buffalo, NY, USA) by direct injection into the aortic cannula (added in 0.4 μL increments over 8 min, i.e. diluted in 26.4 mL of saline) to enable optical mapping.

Hearts were then transferred to another Langendorff system and perfused between 2 and 5 mL per minute, adjusted to ensure a coronary perfusion input pressure of 80 mmHg. Hearts were positioned in a water-jacketed bath (at 37°C) and two spring-loaded Ag/AgCl pellet electrodes (73–0200, Harvard Apparatus, Holliston, MA, USA) placed upon the cardiac surface to provide a surface electrocardiogram (pseudo-ECG). ECG, perfusion pressure, and camera exposure were recorded using AcqKnowledge with a BIOPAC recording system (BIOPAC MP150, BIOPAC Systems, Goleta, CA, USA). Subsequent data analysis was performed using Matlab (Mathworks, Natick, MA, USA).

Hearts were illuminated by a band-pass filtered (ZET642/20X, Chroma Technology, Bellows Falls, VT, USA) red LED (CBT-90R, Luminus, Sunnyvale, CA, USA) and imaged with a Scientific CMOS device (Andor Zyla 5.5, Oxford Instruments, Abingdon, UK) via an MVX10 macro zoom microscope (Olympus Life Science, Tokyo, Japan) and an emission filter (ET585/50M+800/200M, Chroma Technology, Bellows Falls, VT, USA). The camera was controlled, and signals were acquired, using Andor Solis software (Oxford Instruments, Abingdon, UK). Optical mapping data were analyzed using a custom-designed Matlab program (available from the authors on request).

Hearts were positioned in the bath with the ventral side up so that at least part of the epicardial surface of all four chambers was facing the camera used for optical mapping. Baseline ECG and optical mapping recordings were acquired, and hearts were left to recover to a sinus rhythm of 5–6 Hz for at least 20 min. Hearts were then exposed to increasing concentrations of carbachol (3.125, 12.5, 50, 100, 200, 400, 800, and 1600 nM, for 10 min at each concentration) or to isoproterenol (0.3, 1, 3, 10 nM, for 15 min at each concentration). ECG and optical signals were obtained for at least one time point at the end of exposure to each concentration of each drug, with additional recordings when an increase or reduction in spontaneous beating frequency, or an abnormality in conduction or an arrhythmia were detected. If the spontaneous heart rate during exposure to any drug concentration was below 8 Hz, the heart was transiently electrically paced (at 8 Hz), using a concentric electrical stimulation electrode (PI-SNE-100, World Precision Instruments, Sarasota, FL, USA) placed on the apex, while optical mapping recordings were obtained for assessment of apparent ventricular conduction velocity.

Conduction velocity was analyzed using a custom Matlab program (available from the authors on request). The optical mapping data were loaded, normalized on a pixel-by-pixel basis, and inverted. The ventricles were segmented before applying a 5×5 Gaussian spatial filter with a sigma of 0.5. The image was then binned with a 2×2 mask size using bilinear interpolation. A 20×20-pixel region of interest was selected in the center of the field of view of ventricular myocardium, and the data were averaged across multiple paced excitations to reduce noise. An activation map was generated from the resulting optical mapping data, with local activation defined as the time of maximum action potential upstroke velocity. Conduction velocity on the ventricular surface was then calculated using the Bayly method with the resulting velocities, in pixels/second, converted to metric units using a measured scaling factor (40 μm/pixel edge)[18].

### 2.6 Single-cell and single-nucleus RNA-sequencing datasets

Expression of *Sox10* and *S100B* in mouse and human hearts was investigated using single-cell RNA-sequencing (RNA-seq) data [19] or single-nucleus RNA-seq data [20] from previous studies.

### 2.7 Graphic representation and statistics

Prism 9.5.1 software (GraphPad, San Diego, CA, USA) was used for statistical analyses, preparation of graphs and curve fitting. Graphs show mean ± standard deviation. Concentration-response curves show heart rate plotted against the decimal logarithm of drug concentrations (non-linear regression fit). Both isoproterenol half maximal effective concentration (EC50) and carbachol half maximal inhibitory concentration (IC50) were obtained from the resulting drug-response curves. Statistical significance was determined using the Mann–Whitney *U* test or one-way ANOVA. A *p*-value < 0.05 was deemed to indicate a statistically significant difference between means.

## 3. Results

### 3.1 There is a lower density of SC in cardiac and sciatic nerves of old compared to young, mice

To determine whether there is an age-related change of neuron-supporting SC, we quantified the number of cells expressing the SC markers SOX10 or S100B and the total number of cells, estimated by the number of nuclei. We found significantly fewer SOX10+ (Fig 1A, A’) and S100B+ (Fig 1B, B’) cells, as well as fewer cells in total (Fig 1A”) per area of nerve tissue in sciatic nerves from old mice, compared to young mice. The cross-sectional area of individual nerves was not significantly different between age groups (Fig 1C). In cardiac tissue obtained from old mice, significantly smaller ratios of SOX10+ (Fig 2A’) and S100B+ (Fig 2B’) nuclei to total cells were found. Additionally, we found that in hearts from young mice, there were fewer SOX10+ SC than S100B+ SC (Fig 2C). Single-cell or single-nucleus (sc/sn)RNA-seq data from mouse (Fig S2A) and human hearts (Fig S2B) confirmed expression of *Sox10/SOX10* and *S100b/S100B* specifically in nerves (the test did not resolve the neuronal and SC compartments). We could not detect S100B protein in CD31+ cells (endothelial/endocardial cells), Tcf21+ cells (epicardial cells, pericytes, cardiac fibroblasts), or CD68+ cells (macrophages) (Fig S2C, D). Finally, we confirmed, based on cell shape, that S100B+ cells (thin, spindle-shaped cells) are distinct from CM (brick-shaped cells). These results show S100B+ as a suitable marker for cardiac SC and suggest the presence of a smaller number of SC both in sciatic and in cardiac nerves from old mice, compared to young mice.

**Figure 1.**
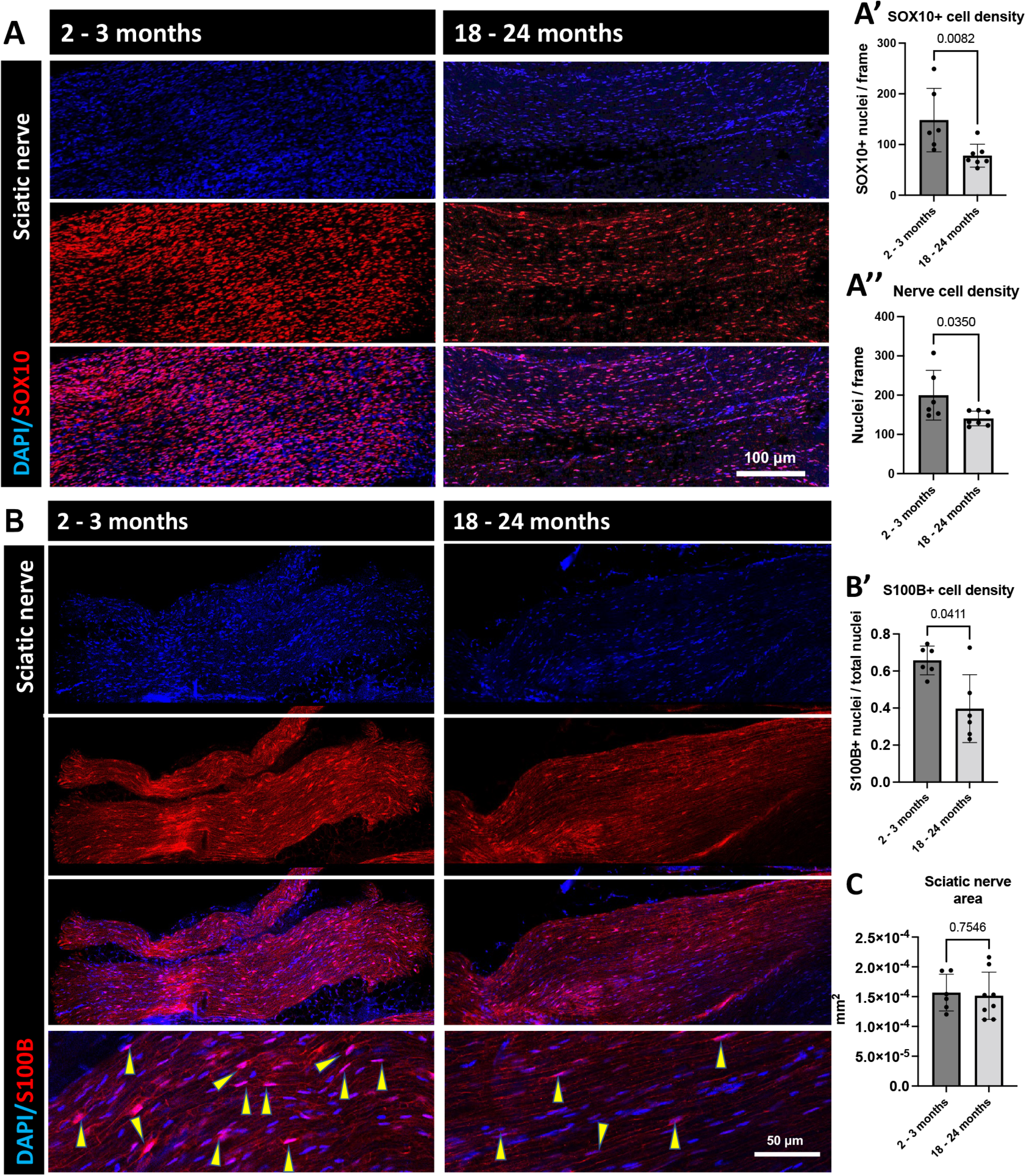
SC density is lower in sciatic nerves isolated from old compared to young mice. (A, B) Representative whole-mount immunofluorescence images show fewer nuclei (detected by DAPI) in general, and fewer SC (detected by SOX10 (A) or S100B (B) co-labelling) in particular. Arrowheads point to S100B+ cells. (A’, B’) Quantification of the spatial density of SOX10+ nuclei (A’, N = 6 [3 (F)emale, 3 (M)ale] from young mice, 7 [4F, 3M] from old mice) or S100B+ nuclei (B’, N = 6 [1F, 5M] young, 6 [6M] old). (A”) Quantification of the spatial density of overall nuclei (N = 6 [3F, 3M] young, 7 [4F, 3M] old). (C) Cross-sectional area measurements of sciatic nerves at their widest position, established by z-stack imaging, from young and old mice (N = 6 [3F, 3M] young, 8 [4F, 4M] old). P value was assessed with the Mann-Whitney *U* test unless stated otherwise.

**Figure 2.**
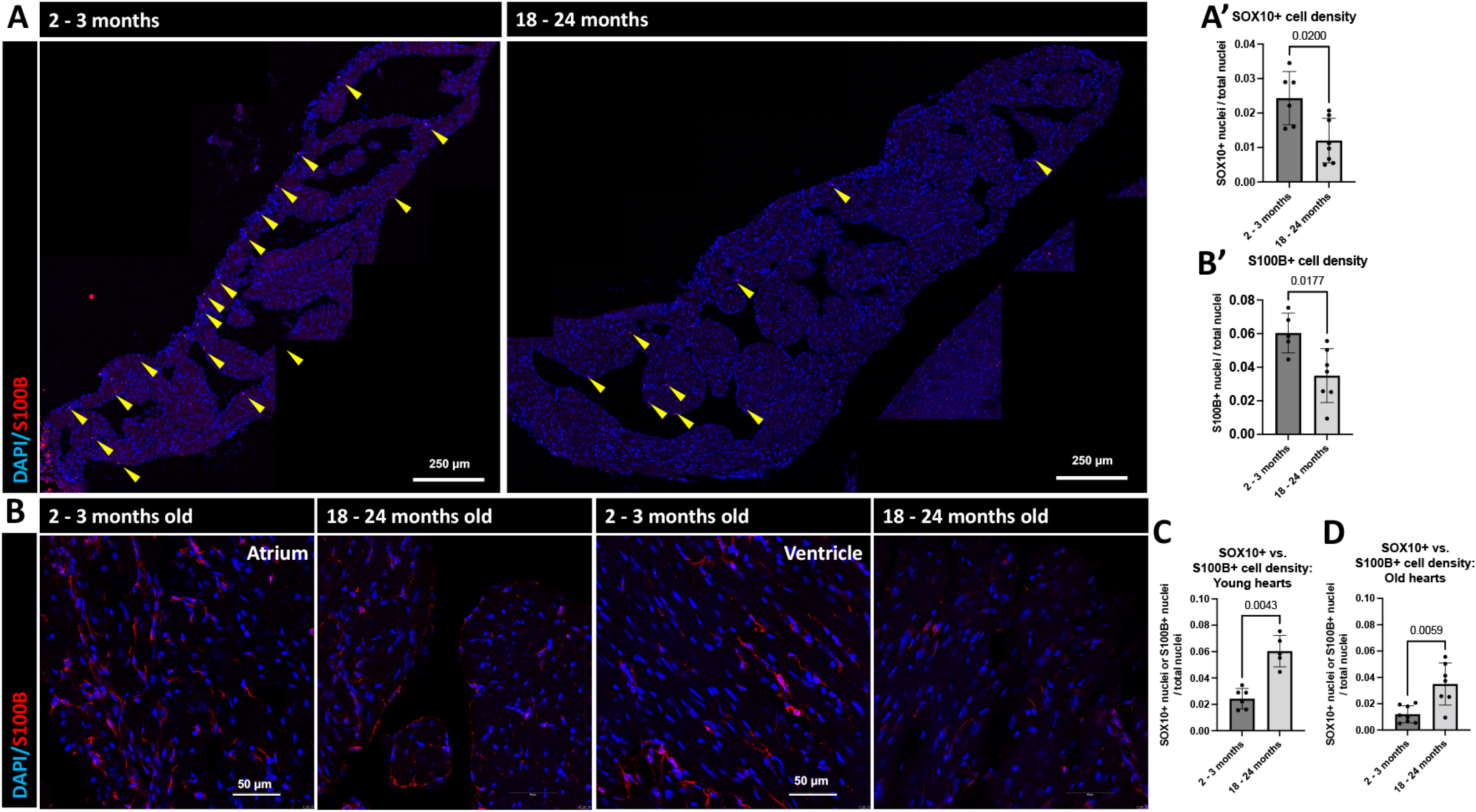
SC density is lower in hearts from old compared to young mice. (A) Representative immunofluorescence of SC nuclei detected by co-staining for DAPI and SOX10 (A) or DAPI and S100B (B). Arrowheads point to SOX10+ nuclei. (A’, B’) Quantification of the spatial density of Sox10+ (A’, N = 6 [3F, 3M] young, 8 [4F, 4M] old) or S100B (B’, N = 5 [3F, 2M] young, 7 [3F, 4M] old) nuclei across the heart. Smaller fractions of SOX10+ and S100B+ nuclei were detected in hearts from old compared to young mice. (C, D) Ratio of SOX10 and S100B nuclei over total nuclei in hearts from young (C, N = 6 [3F, 3M] for SOX10, 5 [3F, 2M] for S100B) and old (D, N = 8 [4F, 4M] for SOX10, 7 [3F, 4M] for S100B) mice. There were substantially fewer SOX10+ nuclei than S100B+ nuclei in hearts from both young and old mice.

### 3.2 SC from old mice show lower proliferative activity and, in the sciatic nerve, higher incidence of apoptosis

To assess whether the fraction of SC that undergo apoptosis differs with age in nerves of the PNS and the iCNS in young and old mice, we performed TUNEL staining of sciatic nerves and hearts, co-stained with SOX10, which is detected in the nuclei, and made the counting of co-localizated TUNEL nuclear signal more precise than S100B. We found a higher fraction of apoptotic cells in general, and a higher fraction of apoptotic SC in particular, in sciatic nerves from old mice, compared to young animals (Fig 3A, A’, A”). However, we did not find statistically significant differences in the heart (Fig 3B, B’, B”). To study the proliferative activity of SC of the PNS and the iCNS, we co-stained tissue with the G2-M marker phospho-histone3 (PH3) and SOX10, and found a substantially lower fraction of PH3+ cells in general in sciatic nerves from old mice, compared to young animals (Fig. 3C”). While the overall fraction of cell-cyling nuclei in the heart was not significantly different between age groups (Fig. 3D”), there was a substantially smaller fraction of PH3+ cells among SC, both in sciatic nerve (Fig 3C, C’) and heart (Fig 3D, D’) of old, compared to young mice. Our data suggest that a reduced rate of proliferation (and an increased rate of cell death in sciatic SC) may underlie the observed age-related differences in SC numbers.

**Figure 3.**
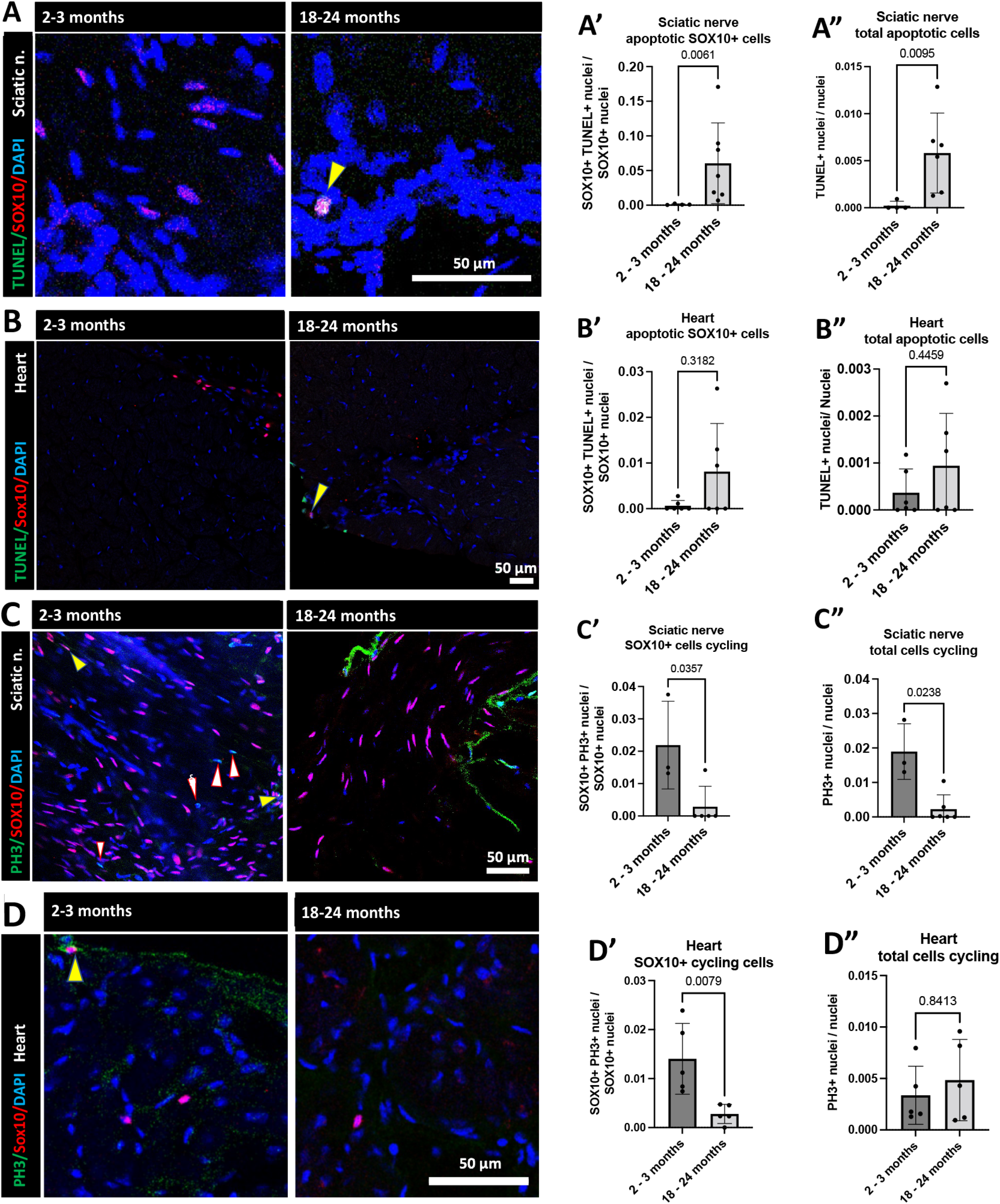
A smaller fraction of SC proliferate in cardiac and sciatic nerves and a larger fraction of SC are apoptotic in sciatic nerve from old compared to young mice. (A, C) Representative immunofluorescence images of sciatic nerve samples show increased SC cell death by TUNEL/SOX10 co-expression, and reduced proliferation in SC by co-localization of SOX10+ nuclei with the G2-M phase marker PH3 in old compared to young mice. Yellow arrowheads point to TUNEL+/SOX10+ or PH3/SOX10+ nuclei. White arrowheads point to PH3+/SOX10-nuclei (A’, C’) Quantification of the number of TUNEL+/SOX10+ nuclei (A’, N= 4 [2F, 2M] young, 7 [3F, 4M] old), and PH3+/SOX10+ nuclei (C’ = 3 [2F, 1M] young, 6 [3F, 3M] old) relative to the total number SOX10+ nuclei. A larger fraction of TUNEL+/SOX10+ nuclei and a smaller fraction of PH3/SOX10 nuclei were detected in sciatic nerves from old compared to young mice. (A”, C”) Quantification of the number of TUNEL+ (A”, N = 4 [2F, 2M] young, 7 [3F, 4M] old) or PH3+ (C”, N = 3 [2F, 1M] young, 6 [3F, 3M] old) nuclei relative to the total number of nuclei. A larger fraction of TUNEL+ and smaller fraction of PH3+ nuclei were detected in the sciatic nerves from old compared to young mice. (B, D) Representative immunofluorescence images of heart samples show TUNEL/SOX10 co-expression for apoptosis, and co-localization of SOX10+ nuclei and PH3 for proliferation assessment in old and young mice. Yellow arrowheads point to TUNEL+/SOX10+ or PH3/SOX10+ nuclei. (B’, D’) Quantification of the number of TUNEL+/SOX10+ nuclei (B’ = 6 [2F, 4M] young, 6 [3F, 3M] old), and PH3+/SOX10+ nuclei (D’ = 5 [2F, 3M] young, 5 [1F, 4M] old) relative to the total number SOX10+ nuclei. No statistically significant difference in the fraction of TUNEL+/SOX10+ nuclei was observed between old and young animals. A lower fraction of PH3+/SOX10+ nuclei was detected in hearts from old compared to young mice. (B”, D”) Quantification of the number of TUNEL+ (B”, N = 6 [2F, 4M] young, 6 [2F, 4M] old) or PH3+ (D”, N = 5 [2F, 3M] young, 5 [1F, 4M] old) nuclei relative to the total number of nuclei. No significant differences in the fraction of apoptotic or proliferating cells overall were detected between hearts from old and young animals.

### 3.3 Parasympathetic nerve fibers are less dense in sciatic and cardiac nerves of old compared to young mice

We performed whole-mount immunofluorescence of sciatic and cardiac nerves from old and young mice. To differentiate and quantify individual nerve fibers, we co-stained with WGA (stains basement membranes). In nerves from the PNS (Fig 4A, A’) and iCNS (Fig 4B, B’), we found significantly lower numbers of parasympathetic nerve fibers (detected by staining against choline acetyltransferase [ChAT]) in old mice compared to young mice. The number of sympathetic nerve fibers (detected by staining against tyrosine hydroxylase [TH]) was lower in sciatic nerves from old mice versus those from young mice, while no significant differences were seen in cardiac sympathetic nerves (Fig 4A”, B”). Our findings show that nerves from old mice have fewer parasympathetic nerve fibers in the PNS and the iCNS, when compared to nerves from young mice.

**Figure 4.**
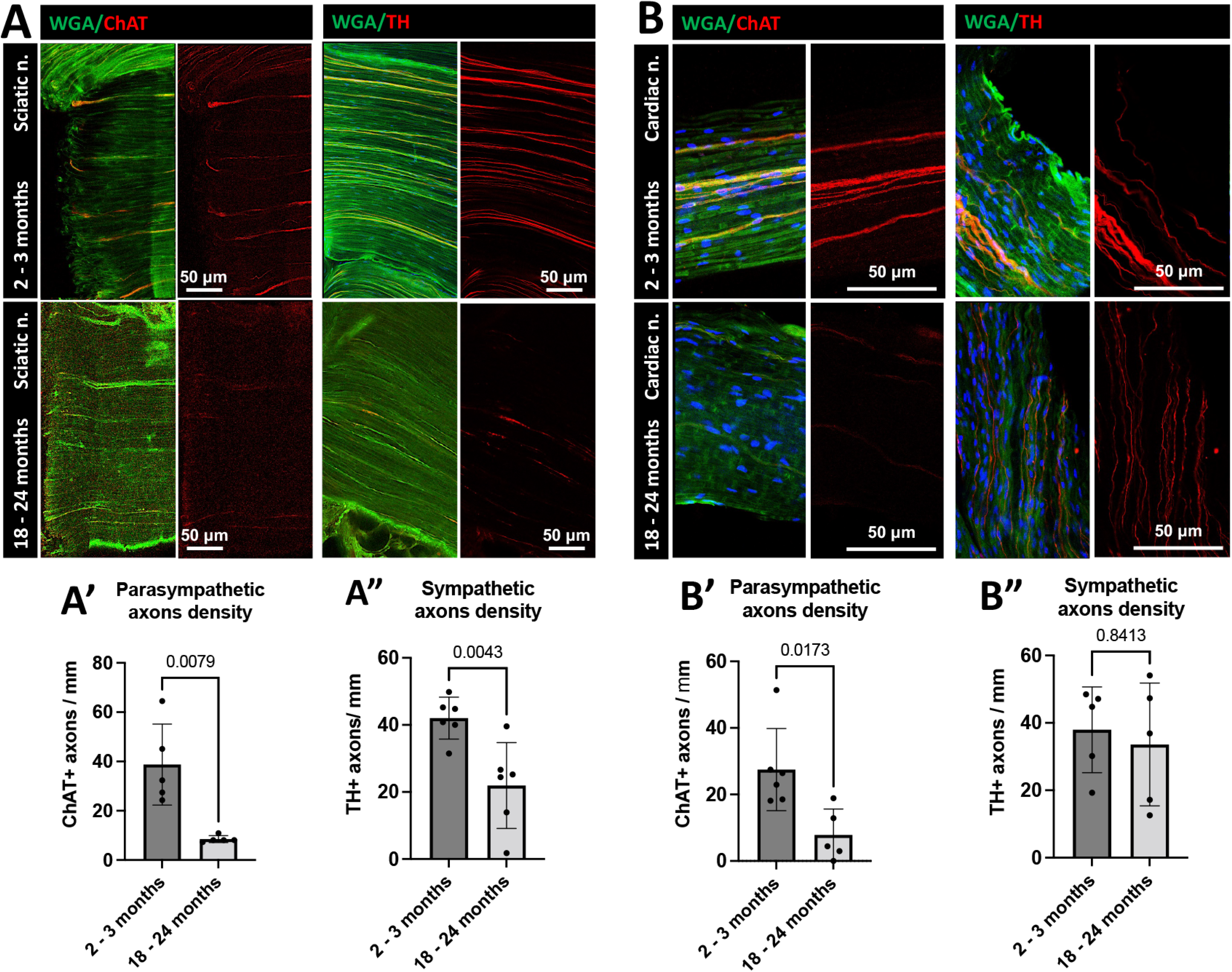
Parasympathetic nerve fiber density in sciatic and cardiac nerves is lower in old compared to young mice. (A, B) Representative whole-mount immunofluorescence images of parasympathetic (ChAT+) and sympathetic (TH+) nerve fibers in sciatic (A) and cardiac (B) nerves from old and young mice. (A’, B’) Quantification of the number of ChAT+ nerve fibers relative to thickness of the sciatic (A’) or cardiac (B’) nerves at the quantification point (widest part of the nerve, as established by z-stack imaging). A lower density of ChAT+ fibers was detected in the sciatic nerves (N = 5 [2F, 3M] young, 5 [2F, 3M] old) and cardiac (N = 5 [3F, 2M] young, 6 [2F, 4M] old) from old compared to young mice. (A”, B”) Quantification of the number of TH+ nerve fibers relative to the thickness of the sciatic (A”) or cardiac (B”) nerve at the quantification point. A lower density of TH+ fibers was detected in the sciatic (A” N = 6 [1F, 5M] young, 6 [1F, 5M] old) but not in the cardiac (B”, N = 5 [3F, 2M] young, 5 [2F, 3M] old) nerves from old compared to young mice.

### 3.4 Acidic polysaccharides and/or glycosaminoglycans are found in nerves from old but not from young mice

As the cross-sectional area of sciatic nerves did not differ between old and young mice (Fig 1C), we explored what might fill the space formerly occupied by SC and nerve fibers. As aging is associated with the development of fibrosis, sclerosis and calcification of soft tissues [21, 22], we assessed cartilage and bone neoformation by histological staining. Staining with Acian blue indicated deposition of acidic polysaccharides and glycosaminoglycans (like proteoglycans, Fig 5), which are involved in cartilage formation, in sciatic nerves from old but not from young mice. No Orange G-stainable deposits were detected, indicating a lack of differentiated bone tissue. The proteoglycan CSPG4 is involved in nerve remodeling and therefore we interrogated whether there were differences in deposits in the sciatic nerve from old and young mice. Immunofluorescence studies showed a higher amount of CSPG4 in sciatic nerve from old mice. On the other hand, no significant differences in CSPG4 abundance were found in TUBB3+ cardiac nerves from old and young mice. Our data point to an accumulation of cartilage-like connective tissue, specifically of CSPG4 in sciatic, but not in cardiac, nerves from old mice.

**Figure 5.**
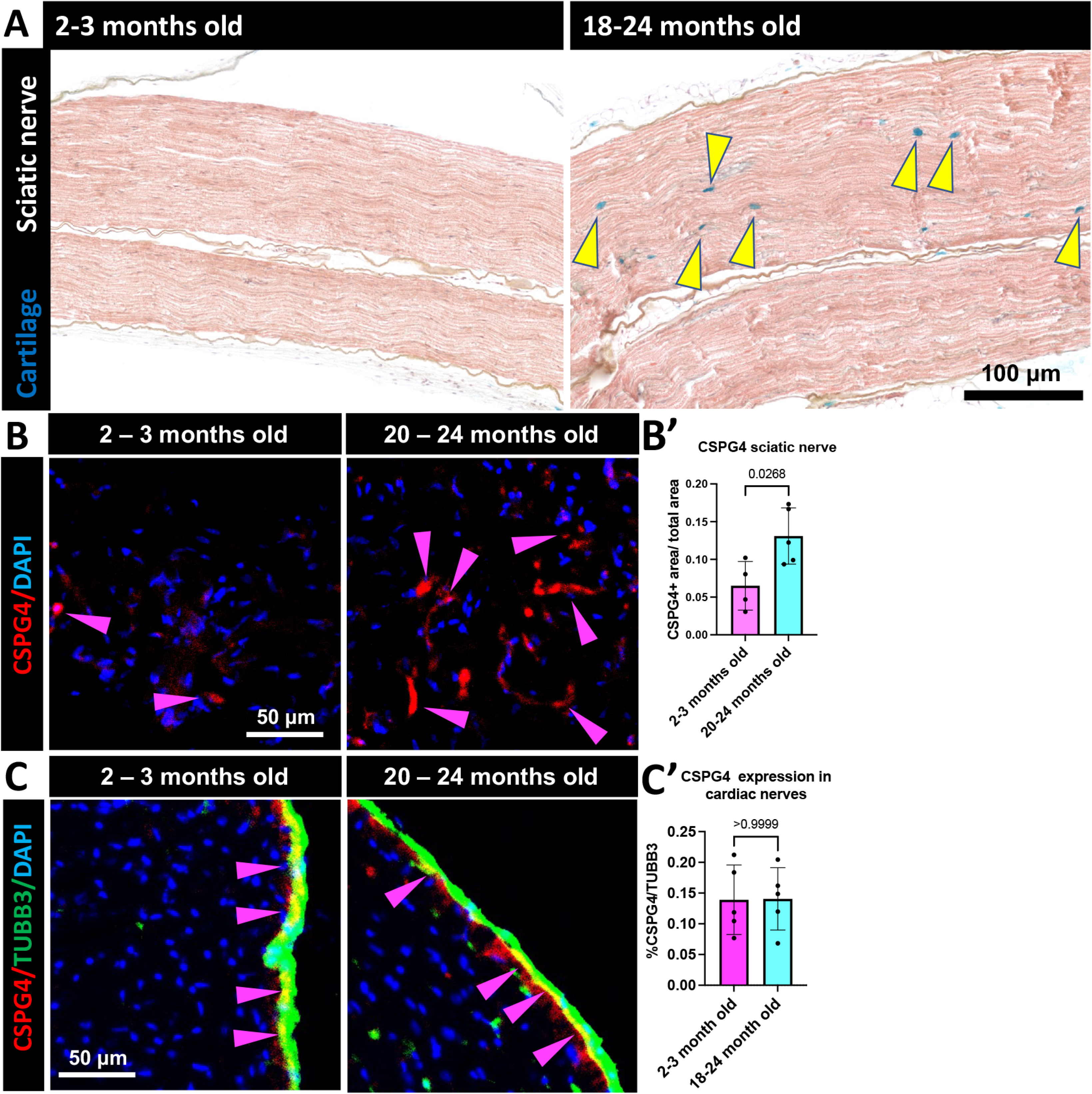
Deposits of CSPG4 can be found in sciatic nerves, but not in cardiac nerves, from old but not young mice. (A) Representative histological images with Alcian Blue/Hematoxylin/Orange G/Eosin staining. In the nerves of old mice, blue deposits that correspond to acidic polysaccharides can be observed (yellow arrowheads); these were not detected in nerves from young mice (N = 5 [2F, 3M] young, 5 [2F, 3M] old). (B) Representative immunofluorescence images of CSPG4 in sciatic nerves from old and young mice. (B’) Quantification of the total area of CSPG4 levels were higher in the sciatic nerves. Significantly more CSPG4 was found in the nerves from old mice, compared to young mice (N = 4 [2F, 2M] young, 5 [3F, 2M] old). (C) Representative immunofluorescence images of the heart from old and young mice do not show significant differences in expression levels of CSPG4 in cardiac nerves detected by TUBB3. (C’) Quantification of the area of CSPG4 that colocalized with TUBB3 (N=5 [3F, 2M] young, 5 [2F, 3M] old).

### 3.5 The expression of M2R in ventricular CM and conduction cells, but not in SAN and AVN cells, is lower in hearts from old compared to young mice

We explored the cardiac localization and expression levels of M2R, the principal receptor for acetylcholine in CM. We did not find significant differences in M2R immunofluorescence signals in pacemaker cells within whole-mount SAN (Fig 6A, A’) tissue and AVN tissue slices (Fig 6C, C’) of young and old mice. Of note, areas surrounding the SAN contained CM with high levels of M2R-fluorescence signal (Fig 6B). On the other hand, we found that the ventricular conduction system (VCS) in hearts from old mice had lower fluorescent density of M2R than the VCS in hearts from young mice (Fig 6C, C”). Finally, we used *MHC^Cre^;R26^ChR2:eYFP^* mice, whose CM express eYFP in their plasma membranes, to study the co-expression of M2R in the cell membrane of ventricular CM. In agreement with our findings in the VCS, M2R fluorescence density was lower in ventricular CM from old compared to young mice (Fig 6D, D’). To corroborate our imaging data, we compared the RNA expression of *M2r/M2R* in ventricular CM of 3 months and 24 months old mice using scRNAseq, and in 40-45 years compared to 70-75 years humans via sn/scRNAseq data (Fig S3). The fraction of CM expressing high levels of M2R was higher in the younger cohort of both species, compared to older samples.

**Figure 6.**
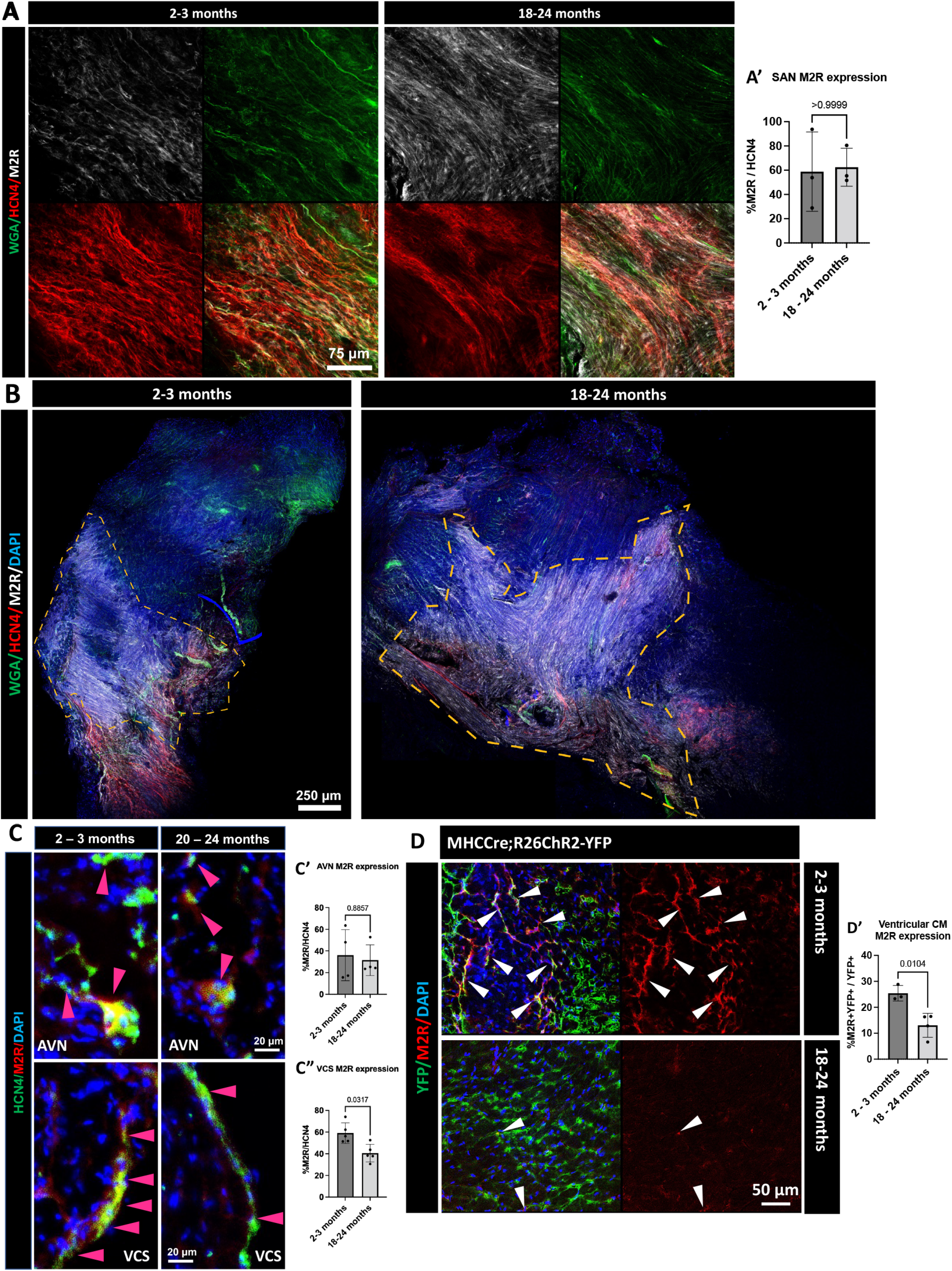
M2R expression is lower in the membrane of ventricular CM of old compared to young mice. (A) Representative whole-mount immunofluorescence images of the SAN from old and young mice do not show significant differences in expression levels of M2R in pacemaker cells. (A’) Quantification of the area of M2R that colocalizes with the pacemaker cell marker HCN4. No significant differences were found between pacemaker tissue from old and young hearts (N = 3 [2F, 1M] young, 3 [1F, 2M] old). (B) Representative whole-mount immunofluorescence images of the SAN from young and old mice show a ring of CM with high levels of M2R fluorescence. (C) Representative immunofluorescence images of M2R show no notable age-related differences expression in the AVN, but significantly lower fluorescence in the VCS of old mice. (C’) Quantification of the area of M2R that colocalizes with the pacemaker cell marker HCN4 in AVN: no significant differences were found between old and young hearts (N=5 [3F, 2M] young, 5 [2F, 3M] old). (C”) Quantification of the area of M2R that colocalizes with HCN4 in VCS: significantly lower levels of M2R are seen in old, compared to young mice (N=5 [3F, 2M] young, 5 [2F, 3M] old). (D) Representative immunofluorescence images of ventricular myocardium from old and young *MHC^Cre^;R26^ChR2-YFP^*mice. Hearts from old mice show lower expression of MR2 on the membrane of CM (YFP+) compared to young mice. White arrowheads point to YFP+/M2R+ cell membranes (D’) Quantification of the area of M2R that colocalize with the membrane of CM shows a significantly lower abundance of M2R in the membrane of ventricular CM from old, compared to young mice (N = 3 [1F, 2M] young, 4 [2F, 2M] old).

### 3.6 Basal heart rate is lower and ventricular arrhythmogenesis is more frequent upon carbachol application in Langendorff-perfused hearts from old compared to young mice

We stimulated Langendorff-perfused hearts with increasing concentrations of isoproterenol (β-adrenoreceptor agonist) and carbachol (muscarinic/nicotinic receptor agonist), and recorded heart rate and ventricular conduction properties. While hearts from old mice were significantly heavier than those from young mice, there was no significant difference in the heart weight to body weight ratio (Fig S4). In basal conditions, the hearts of old animals showed significantly lower spontaneous heart rates (Fig. 7A). Heart rate increased when stimulated with increasing concentrations of isoproterenol, with no significant differences between hearts from old and young mice (Fig. 7B, D). Similarly, upon stimulation with increasing concentrations of carbachol, heart rate reduction was not significantly different in old and young mice (Fig. 7C, E). While there was no significant difference in basal PQ intervals of hearts from young and old mice, the PQ interval of hearts from old mice increased upon exposure to 400 nM carbachol, whereas no significant change was seen in hearts from young mice (Fig. 7F, Fig S5). No significant differences were found in action potential duration and apparent conduction velocity, as assessed by optical mapping, of ventricular myocardium in old and young mice, and no significant differences were observed with or without carbachol (Fig. 7H). Nonetheless, hearts of old mice were significantly more prone to develop ventricular arrhythmias (tachycardia and fibrillation; Fig. 7H, I), and they did so at lower carbachol concentrations than hearts of young mice.

**Figure 7.**
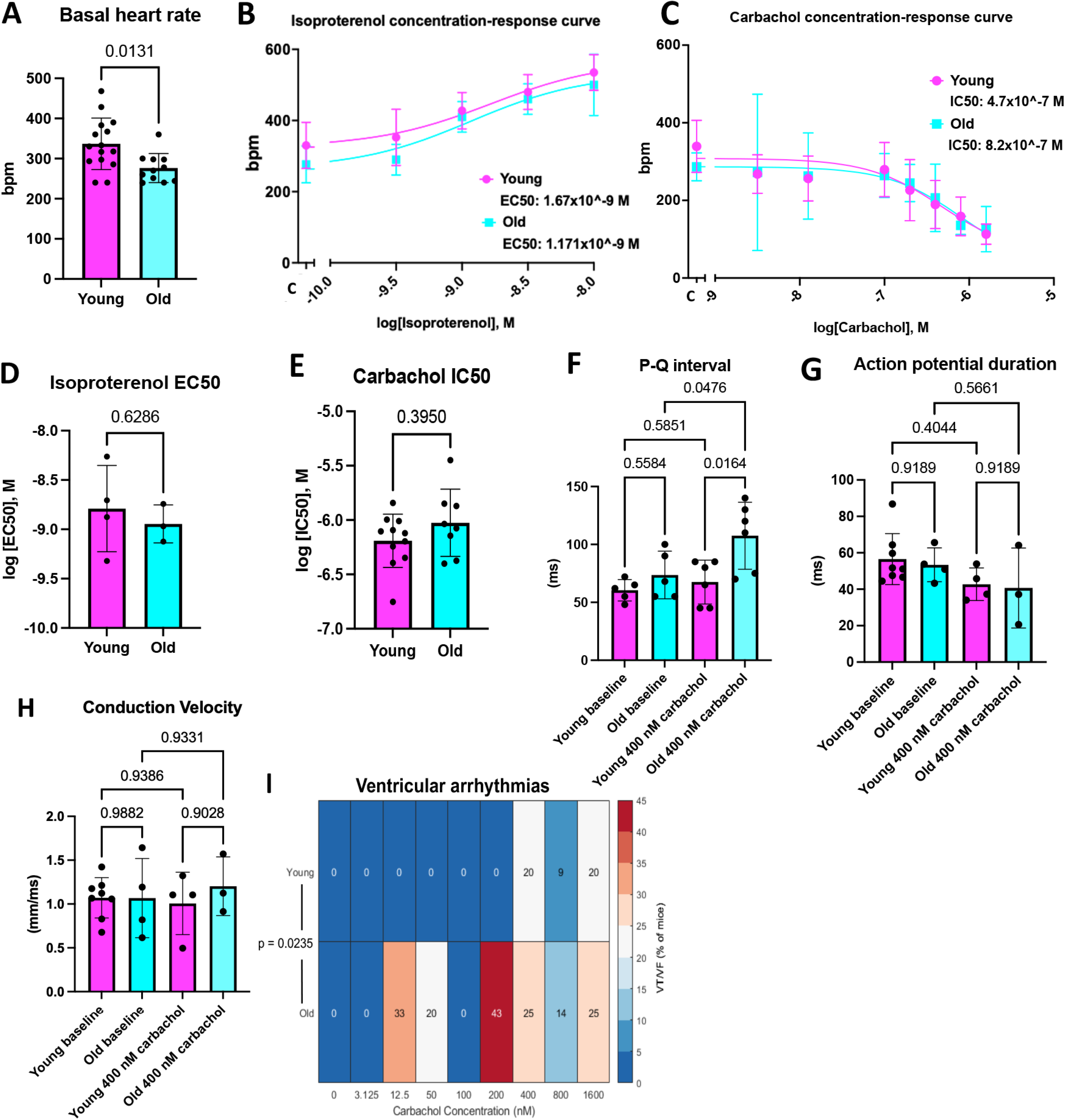
Langendorff-perfused hearts from old mice have a longer PQ interval and are more prone to ventricular arrhythmogenesis upon carbachol stimulation, compared to hearts from young mice. (A) Basal heart rate (beats per minute [bpm]) of Langendorff-perfused hearts from young and old mice (N = 11 [7F, 4M] young, 8 [3F, 5M] old). (B) Concentration-response curve of isoproterenol effects on spontaneous beating rate (in beats per minute; bpm) of Langendorff-perfused hearts from young and old mice. EC50-values from hearts of young animals and old animals are given in the figure (N = 4 [2F, 2M] young, 3 [3M] old). (C) Concentration-response curve of carbachol effects on spontaneous beating rate (in bpm) of Langendorff-perfused hearts from young and old mice. IC50-values from hearts of young animals and old animals are given in the figure (N = 11 [7F, 4M] young, 8 [3F, 5M] old). (D) Isoproterenol EC50 comparison of individual young and old mice. No statistically significance difference was found. EC50 is expressed as the negative log of the molar concentration. (E) Carbachol IC50 comparison of individual young and old mice; no statistically significance difference was found. IC50 is expressed as the negative log of the IC50. (F) PQ interval in Langendorff-perfused hearts from young and old mice before (baseline) and during application of 400 nM carbachol. (N = 4 [2F, 2M] young, 5 [2F, 3M] old). (G) Action potential duration at 80% repolarization in optical mapping data from Langendorff-perfused hearts of young and old mice while paced at 8 Hz, prior to (baseline, N = 8 young [3F, 5M], 4 old [2F, 2M]) and during administration of 400 nM carbachol (N = 4 young [3F, 1M], 3 old [2F, 1M). (H) Apparent ventricular conduction velocity in Langendorff-perfused hearts from young and old mice while paced at 8 Hz, prior to (baseline, (N = 8 young [3F, 5M], 4 old [2F, 2M]) and during administration of 400 nM carbachol (N = 4 young [3F, 1M], 3 old [2F, 1M]). (I) Percentage of mice exhibiting spontaneous ventricular tachycardia (VT) or ventricular fibrillation (VF) in response to exposure to increasing carbachol concentrations. Hearts from old mice were significantly more likely to exhibit VT/VF when exposed to carbachol, and exhibited arrhythmias at carbachol concentrations that are lower than both the age-specific IC50, compared to young mice. Statistical significance was assessed by one-way ANOVA.

## 4. Discussion

We examined age-associated differences of PNS and iCNS and found that in comparison to young (2-3 months) mice, that old mice (20-24 months) are characterized by: (1) lower densities of SC and nerve fibers in cardiac and sciatic nerves, (2) presence of cartilaginous ECM deposits in sciatic nerves, and (3) a higher susceptibility to the development of ventricular arrhythmias when exposed to carbachol.

A reliable marker protein for cardiac SC is a prerequisite for the immunohistochemical study of the distribution and density of SC. We have previously used SOX10 to identify cardiac SC [3], but this marker is known to be related to embryonic and postnatal development and could therefore be unsuitable for identification of mature SC [23]. Here, we observe that the marker proteins SOX10 and S100B are co-expressed by myelinating and Remak SC in the PNS. Both proteins are also specifically expressed in SC of the iCNS, though the number of SOX10+ SC was significantly lower than the number of S100B+ SC in the heart. Given the important role of SOX10 in SC differentiation/dedifferentiation and proliferation, it may be expressed in a subpopulation of more plastic or even “ready-to-heal” cardiac SC in hearts from adult mice [24–27]. Exploring differences in expression patterns and possible links to cell function of SOX10+ and S100B+ cardiac SC forms an interesting target for future work. Our data suggest that S100B is a pan-cardiac SC marker, while SOX10 labels a subset of cardiac SC only.

Our data demonstrate an age-related reduction in SC density in peripheral nerves and the iCNS. With exception of the heart, many organs, including the liver, thymus and skeletal muscle, are known to lose mass during aging [28]. An increase in the rate of programmed cell death has also been associated with aging in different tissues, including the brain, skeletal muscle and the heart [1, 29, 30]. Here, an age-related increase in the fraction of apoptotic SC was seen in sciatic nerve, but not the heart. The observed reduction in SC density in hearts of old mice may thus not be explained by higher apoptosis rates, at least not at the time of observation. In addition to apoptosis, aging has been linked with a reduced rate of proliferation [1]. A substantially lower reduction in the fraction of proliferating SC was observed both in sciatic nerves and in the heart of old mice, which could contribute to the reduction in SC numbers during aging and to the previously described loss of neural regenerative capacity in aged individuals [31]. In conclusion, aging is associated with a reduced density of SC in PNS and iCNS of mice, which may be linked to a loss of the proliferative capacity of SC.

SC maintain axonal, synaptic and gap junction homeostasis, and dysfunction of SC is associated with neuromuscular junction abnormalities and nerve fiber degeneration. Here, we confirm an aging-related loss of sympathetic and parasympathetic nerve fibers in the PNS [31], while in the iCNS only parasympathetic nerve fibers appear to be reduced. This data, along with the lower M2R expression levels on the membrane of ventricular CM from old hearts, which has also been shown in human right atrial tissue by radioligand binding studies [32] supports the idea of an age-related attenuation of parasympathetic control of the heart, potentially leading to an imbalance in sympathetic *versus* parasympathetic stimulation. As laboratory mice have a life-span of up to 30 months, it is possible, of course, that cardiac sympathetic nerves are lost at a later stage in their life than the here assessed 20-24 months of age.

A reduction in cell numbers without a concurrent change in cross-sectional area of nerves raises the question of what matter might fill the resulting space. Some soft tissues, such as blood vessels or cardiac valves, calcify with age [21, 22]. We found neo-cartilage ECM deposits (proteoglycans, for example CSPG4) in sciatic nerves from old but not young mice, but we did not detect fully formed bone tissue. Proteoglycans are known to serve as “stop/go” signals during nerve development and they have been reported to prevent cardiac reinnervation after myocardial infarction [33]. Interestingly, CSPG4 deposits were not higher in cardiac nerves from old, compared to young mice. We found no age-related differences in sympathetic nerve fiber density either, while in the sciatic nerves this density was already low. A possible explanation for these observations is that the cardiac nerves degenerate later in life, than musculoskeletal nerves. Alternatively, differences in structural complexity of the nerves (sciatic nerves have a proper epineurium, while nerves inside the cardiac muscle are much simpler, down to single axons) could mean that cells responsible for CSPG4 deposition are not present in cardiac nerves. The question of whether the accumulation of proteoglycans in nerves from old mice might drive axonal repulsion, or whether proteoglycan deposition is a consequence of nerve fiber loss, has not been addressed. In summary, we found that cardiac nerves from old mice exhibit fewer parasympathetic nerve fibers and that sciatic nerves from old mice are characterized by lower pan-axonal density, and increased cartilaginous proteoglycan, like CSPG4, deposits, compared to young mice.

While our work provides novel insights into structural differences of PNS and iCNS in young vs old mice, there are certain technical limitations. In our analysis, we compared the number of SC to the total number of nuclei in the tissue per area of tissue. However, while cells of the PNS are mononucleated, a large proportion of murine cardiomyocytes are binucleated [34]. Therefore, (i) the fraction of cardiac SC may be higher than reflected by our nucleus staining [35], and (ii) this scaling may differ between mice of different age if the share of multi-nucleated CM increases between the ages of 2-3 and 20-24 months. Additionally, due to the large intra-individual variability in cardiac nerve thickness (as opposed to sciatic nerves), we were unable to provide a quantification of cardiac aging-related dynamics in iCNS nerve cross-sectional area.

We further assessed functional alterations associated with age-related differences in iCNS structure. In our *ex vivo* studies, we found lower basal heart rates in hearts from old compared to young mice. This finding agrees with previous reports in mice [36], but differs from observations in humans, where basal heart rate does not change with age in the adult [37], although an age-associated slowdown of SAN automaticity has been also reported not only in humans, but also dogs and rabbits [38, 39, 40]. Recently, a prolonged PQ interval has been described in aged rats when compared with younger counterparts [41]. While this matches our data, only a prolonged PR has been reported in humans [42]. Isoproterenol increased SAN rate in hearts from both young and old mice, and we did not detect a significant difference in the sensitivity of the SAN (based on EC50) to isoproterenol between age groups, contrary to previous studies that showed a lower sensitivity of beating rate to isoproterenol in hearts from old mice [43]. The fact that we used concentrations of isoproterenol that were four orders of magnitude lower than those used in previous studies may provide a possible explanation for these conflicting results. When considering that a change from 0.44 nM to 2.7 nM in plasma epinephrine is increases heart rate by 17 bpm in humans [44], and that the concentration of epinephrine provided as an emergency intervention in cardiac arrest is ∼250 nM (given approximately 5 liters of blood and 15 liters of extracellular liquid), we consider that our dosage range as physiologically relevant.

In hearts from both age groups, administration of increasing concentrations of carbachol reduced heart rate. As with isoproterenol application, heart rate sensitivity to carbachol (based on IC50) did not differ significantly in hearts from old (based on IC50) compared to young mice. Interestingly, we found a higher incidence of spontaneous ventricular tachycardia and fibrillation in hearts from old mice upon exposure to increasing carbachol concentrations. Arrhythmogenesis started to manifest itself at carbachol concentrations 30 times lower (from 12.5 nM) than the IC50 for heart rate responses (820 nM). Previous work in young mice (2.5 to 4 months) had shown that parasympathetic stimulation favors ventricular arrhythmogenesis in response to burst pacing [45]. Spontaneous ventricular arrhythmias also occurred upon carbachol application in a LQT3 disease mouse model [46]. Our results differ from observation in humans of advanced age, where increased parasympathetic input is more likely to be associated with atrial, rather than ventricular arrhythmias (with the exception of Brugada’s syndrome). Apart from species-related differences (we used mouse models with a one order of magnitude higher intrinsic heart rates), we used an *ex vivo* Langendorff-perfused model, where the systemic balance of central innervation is absent. This allowed us to isolate effects of pharmacological stimulation, whose molecular mechanism require further investigation - given that an age-dependent increase in carbachol-induced arrhythmogenesis has not been reported previously.

It is plausible, of course, that hearts from old mice are more prone to arrhythmia simply due (i) to their increased size (to the point that the ventricular dimensions may exceed the critical wavelength for reentry, thereby promoting sustainable ventricular arrhythmias [47, 48]) and/or (ii) to a higher degree of tissue fibrosis. In this context, carbachol may sensitize Langendorff-perfused hearts to arrhythmia formation as previously shown [46, 49]. Our results may be regarded as hypothesis-forming, as it is possible that the combination of increased heart size and/or fibrosis with putative changes in the sensitivity of the ventricular cardiac conduction system to carbachol (as indicated by decreased density of M2R), may promote the occurrence of arrhythmogenic triggers, perhaps in lower order ventricular pacemaker cells. To assess this hypothesis, further research is required.

In conclusion, our data show previously unknown features of PNS and iCNS aging, including a loss of SC and relative reduction in parasympathetic nerve fibers, accompanied by a reduction in M2R levels in ventricular CM and ventricular conduction system cells. Additionally, they pose new questions for follow-up research, e.g. regarding contributions of the iCNS and of PNS receptor presence and distribution on ventricular arrhythmogenesis in aged hearts, such as observed *ex vivo* upon parasympathetic stimulation.

## Supporting information

Saasu_E, Supplementary info

## Acknowledgements

We thank, Dr. Katherine E. Yutzey, Dr. Crystal M Ripplinger, Dr. Beth A. Habecker and Dr. Achim Lother for their scientific advice. We also thank the staff of the IEKM and SFB1425 for valuable feedback. We acknowledge the microscopy facility SCI-MED (Super-Resolution Confocal/Multiphoton Imaging for Multiparametric Experimental Designs) at IEKM, Freiburg, for providing expertise and access to imaging setups and analysis workstations.

## Author Contributions

Study design was done by CZJ, FSW and LH. Experimental procedures were executed by ES, GT, CZJ, PI, MCF, TB and LH. Data acquisition, analysis, and interpretation were carried out by ES, GT, DS, CZJ, JM, MK, MCF, TB, BC, SP, UR, PK and LH. Figures and manuscript were prepared by CZJ, FSW, PK and LH. The final version of the manuscript was approved by all authors.

## Abbreviation list

AVN: Atrio-ventricular node
ChAT: Choline acetyltransferase
CM: Cardiomyocyte
CSPG4: Chondroitin sulfate proteoglycan 4
DAPI: 4′,6-Diamidino-2-phenylindole
EC50: Half maximal effective concentration
eYFP: Enhanced yellow fluorescent protein
iCNS: Intracardiac nervous system
IC50: Half maximal inhibitory concentration
M2R: Muscarinic acetylcholine receptor M2
OCT: Optimal cutting temperature compound
PBS: Phosphate-buffered saline
PH3: Phospho-histone3
PNS: Peripheral nervous system
RNA-seq: RNA-sequencing
RT: Room temperature
SAN: Sino-atrial node
SC: Schwann cell/s
SOX10: SRY-box transcription factor 10
S100B: S100 calcium-binding protein B
TUBB3: Beta 3 Tubulin
TH: Tyrosine hydroxylase
TUNEL: dUTP nick end labelling
VCS: Ventricular conduction system
WGA: Wheat Germ Aggultinin

## References

[1] C. Lopez-Otin, M.A. Blasco, L. Partridge, M. Serrano, G. Kroemer, The hallmarks of aging, Cell 153 (2013) 1194–217. 10.1016/j.cell.2013.05.039.

[2] M. Gonzalez-Freire, R. de Cabo, S.A. Studenski, L. Ferrucci, The neuromuscular junction: aging at the crossroad between nerves and muscle, Front Aging Neurosci 6 (2014) 208. 10.3389/fnagi.2014.00208

[3] L. Hortells, E.C. Meyer, Z.M. Thomas, K.E. Yutzey, Periostin-expressing Schwann cells and endoneurial cardiac fibroblasts contribute to sympathetic nerve fasciculation after birth, J Mol Cell Cardiol 154 (2021) 124–136. 10.1016/j.yjmcc.2021.02.001

[4] J. Wolbert, X. Li, M. Heming, A.K. Mausberg, D. Akkermann, C. Frydrychowicz, R. Fledrich, L. Groeneweg, C. Schulz, M. Stettner, N. Alonso Gonzalez, H. Wiendl, R. Stassart, G. Meyer Zu Horste, Redefining the heterogeneity of peripheral nerve cells in health and autoimmunity, Proc Natl Acad Sci U S A 117 (2020) 9466–9476. 10.1073/pnas.1912139117

[5] N. Pauziene, D.H. Pauza, R. Stropus, Morphology of human intracardiac nerves: an electron microscope study, J Anat 197 Pt 3 (2000) 437–59. 10.1046/j.1469-7580.2000.19730437.x

[6] J.A. Armour, Cardiac neuronal hierarchy in health and disease, Am J Physiol Regul Integr Comp Physiol 287 (2004) R262–71. 10.1152/ajpregu.00183.2004

[7] L.J. Wagstaff, J.A. Gomez-Sanchez, S.V. Fazal, G.W. Otto, A.M. Kilpatrick, K. Michael, L.Y.N. Wong, K.H. Ma, M. Turmaine, J. Svaren, T. Gordon, P. Arthur-Farraj, S. Velasco-Aviles, H. Cabedo, C. Benito, R. Mirsky, K.R. Jessen, Failures of nerve regeneration caused by aging or chronic denervation are rescued by restoring Schwann cell c-Jun, Elife 10 (2021). 10.7554/eLife.62232

[8] G. Valdez, J.C. Tapia, H. Kang, G.D. Clemenson, Jr., F.H. Gage, J.W. Lichtman, J.R. Sanes, Attenuation of age-related changes in mouse neuromuscular synapses by caloric restriction and exercise, Proc Natl Acad Sci U S A 107 (2010) 14863–8. 10.1073/pnas.1002220107

[9] A. Elia, A. Cannavo, G. Gambino, M. Cimini, N. Ferrara, R. Kishore, N. Paolocci, G. Rengo, Aging is associated with cardiac autonomic nerve fiber depletion and reduced cardiac and circulating BDNF levels, J Geriatr Cardiol 18 (2021) 549–559. 10.11909/j.issn.1671-5411.2021.07.009

[10] D.D. Gibbons, E.M. Southerland, D.B. Hoover, E. Beaumont, J.A. Armour, J.L. Ardell, Neuromodulation targets intrinsic cardiac neurons to attenuate neuronally mediated atrial arrhythmias, Am J Physiol Regul Integr Comp Physiol 302 (2012) R357–64. 10.1152/ajpregu.00535.2011

[11] K. Shivkumar, O.A. Ajijola, I. Anand, J.A. Armour, P.S. Chen, M. Esler, G.M. De Ferrari, M.C. Fishbein, J.J. Goldberger, R.M. Harper, M.J. Joyner, S.S. Khalsa, R. Kumar, R. Lane, A. Mahajan, S. Po, P.J. Schwartz, V.K. Somers, M. Valderrabano, M. Vaseghi, D.P. Zipes, Clinical neurocardiology defining the value of neuroscience-based cardiovascular therapeutics, J Physiol 594 (2016) 3911–54. 10.1113/JP271870

[12] K. Scherschel, K. Hedenus, C. Jungen, M.D. Lemoine, N. Rubsamen, M.W. Veldkamp, N. Klatt, D. Lindner, D. Westermann, S. Casini, P. Kuklik, C. Eickholt, N. Klocker, K. Shivkumar, T. Christ, T. Zeller, S. Willems, C. Meyer, Cardiac glial cells release neurotrophic S100B upon catheter-based treatment of atrial fibrillation, Sci Transl Med 11 (2019). 10.1126/scitranslmed.aav7770.

[13] A. Acharya, S.T. Baek, G. Huang, B. Eskiocak, S. Goetsch, C.Y. Sung, S. Banfi, M.F. Sauer, G.S. Olsen, J.S. Duffield, E.N. Olson, M.D. Tallquist, The bHLH transcription factor Tcf21 is required for lineage-specific EMT of cardiac fibroblast progenitors, Development 139 (2012) 2139–49. 10.1242/dev.079970

[14] K. Rysevaite, I. Saburkina, N. Pauziene, R. Vaitkevicius, S.F. Noujaim, J. Jalife, D.H. Pauza, Immunohistochemical characterization of the intrinsic cardiac neural plexus in whole-mount mouse heart preparations, Heart Rhythm 8 (2011) 731–8. 10.1016/j.hrthm.2011.01.013

[15] K. Rysevaite, I. Saburkina, N. Pauziene, S.F. Noujaim, J. Jalife, D.H. Pauza, Morphologic pattern of the intrinsic ganglionated nerve plexus in mouse heart, Heart Rhythm 8 (2011) 448–54. 10.1016/j.hrthm.2010.11.019

[16] E.A. MacDonald, J. Madl, J. Greiner, A.F. Ramadan, S.M. Wells, A.G. Torrente, P. Kohl, E.A. Rog-Zielinska, T.A. Quinn, Sinoatrial node structure, mechanics, electrophysiology and the chronotropic response to stretch in rabbit and mouse, Front Physiol 11 (2020) 809. 10.3389/fphys.2020.00809

[17] S. Berg, D. Kutra, T. Kroeger, C.N. Straehle, B.X. Kausler, C. Haubold, M. Schiegg, J. Ales, T. Beier, M. Rudy, K. Eren, J.I. Cervantes, B. Xu, F. Beuttenmueller, A. Wolny, C. Zhang, U. Koethe, F.A. Hamprecht, A. Kreshuk, ilastik: interactive machine learning for (bio)image analysis, Nat Methods 16 (2019) 1226–1232. 10.1038/s41592-019-0582-9

[18] P.V. Bayly, B.H. KenKnight, J.M. Rogers, R.E. Hillsley, R.E. Ideker, W.M. Smith, Estimation of conduction velocity vector fields from epicardial mapping data, IEEE Trans Biomed Eng 45 (1998) 563–71. 10.1109/10.668746

[19] C. Tabula Muris, c. Overall, c. Logistical, c. Organ, processing, p. Library, sequencing, a. Computational data, a. Cell type, g. Writing, g. Supplemental text writing, i. Principal, Single-cell transcriptomics of 20 mouse organs creates a Tabula Muris, Nature 562 (2018) 367–372. 10.1038/s41586-018-0590-4

[20] J.D. Hocker, O.B. Poirion, F. Zhu, J. Buchanan, K. Zhang, J. Chiou, T.M. Wang, Q. Zhang, X. Hou, Y.E. Li, Y. Zhang, E.N. Farah, A. Wang, A.D. McCulloch, K.J. Gaulton, B. Ren, N.C. Chi, S. Preissl, Cardiac cell type-specific gene regulatory programs and disease risk association, Sci Adv 7 (2021). 10.1126/sciadv.abf1444

[21] M.V. Gomez-Stallons, J.T. Tretter, K. Hassel, O. Gonzalez-Ramos, D. Amofa, N.J. Ollberding, W. Mazur, J.K. Choo, J.M. Smith, D.J. Kereiakes, K.E. Yutzey, Calcification and extracellular matrix dysregulation in human postmortem and surgical aortic valves, Heart 105 (2019) 1616–1621. 10.1136/heartjnl-2019-314879

[22] L. Hortells, C. Sosa, N. Guillen, S. Lucea, A. Millan, V. Sorribas, Identifying early pathogenic events during vascular calcification in uremic rats, Kidney Int 92 (2017) 1384–1394. 10.1016/j.kint.2017.06.019

[23] K.R. Jessen, R. Mirsky, Schwann Cell Precursors; Multipotent glial cells in embryonic nerves, Front Mol Neurosci 12 (2019) 69. 10.3389/fnmol.2019.00069

[24] J.K. Kim, H.J. Lee, H.T. Park, Two faces of Schwann cell dedifferentiation in peripheral neurodegenerative diseases: pro-demyelinating and axon-preservative functions, Neural Regen Res 9 (2014) 1952–4. 10.4103/1673-5374.145370

[25] J. Kim, L. Lo, E. Dormand, D.J. Anderson, SOX10 maintains multipotency and inhibits neuronal differentiation of neural crest stem cells, Neuron 38 (2003) 17–31. 10.1016/s0896-6273(03)00163-6

[26] S. Britsch, D.E. Goerich, D. Riethmacher, R.I. Peirano, M. Rossner, K.A. Nave, C. Birchmeier, M. Wegner, The transcription factor Sox10 is a key regulator of peripheral glial development, Genes Dev 15 (2001) 66–78. 10.1101/gad.186601

[27] K. Kuhlbrodt, B. Herbarth, E. Sock, I. Hermans-Borgmeyer, M. Wegner, Sox10, a novel transcriptional modulator in glial cells, J Neurosci 18 (1998) 237–50. 10.1523/JNEUROSCI.18-01-00237.1998

[28] Q. He, S. Heshka, J. Albu, L. Boxt, N. Krasnow, M. Elia, D. Gallagher, Smaller organ mass with greater age, except for heart, J Appl Physiol (1985) 106 (2009) 1780–4. 10.1152/japplphysiol.90454.2008

[29] R.B. Richardson, D.S. Allan, Y. Le, Greater organ involution in highly proliferative tissues associated with the early onset and acceleration of ageing in humans, Exp Gerontol 55 (2014) 80–91. 10.1016/j.exger.2014.03.015

[30] M. Pollack, S. Phaneuf, A. Dirks, C. Leeuwenburgh, The role of apoptosis in the normal aging brain, skeletal muscle, and heart, Ann N Y Acad Sci 959 (2002) 93–107. DOI: 10.1111/j.1749-6632.2002.tb02086.x

[31] E. Verdu, D. Ceballos, J.J. Vilches, X. Navarro, Influence of aging on peripheral nerve function and regeneration, J Peripher Nerv Syst 5 (2000) 191–208. 10.1046/j.1529-8027.2000.00026.x.

[32] O.E. Brodde, U. Konschak, K. Becker, F. Ruter, U. Poller, J. Jakubetz, J. Radke, H.R. Zerkowski, Cardiac muscarinic receptors decrease with age. In vitro and in vivo studies, J Clin Invest 101(2) (1998) 471–8. 10.1172/JCI1113.

[33] M.R. Blake, D.C. Parrish, M.A. Staffenson, M.A. Johnson, W.R. Woodward, B.A. Habecker, Loss of chondroitin sulfate proteoglycan sulfation allows delayed sympathetic reinnervation after cardiac ischemia-reperfusion, Physiol Rep 11 (2023) e15702. 10.14814/phy2.15702

[34] N. Velayutham, C.M. Alfieri, E.J. Agnew, K.W. Riggs, R.S. Baker, S.R. Ponny, F. Zafar, K.E. Yutzey, Cardiomyocyte cell cycling, maturation, and growth by multinucleation in postnatal swine, J Mol Cell Cardiol 146 (2020) 95–108.

[35] N. Anto Michel, S. Ljubojevic-Holzer, H. Bugger, A. Zirlik, Cellular Heterogeneity of the heart, Front Cardiovasc Med 9 (2022) 868466. 10.1016/j.yjmcc.2020.07.004

[36] C. Piantoni, L. Carnevali, D. Molla, A. Barbuti, D. DiFrancesco, A. Bucchi, M. Baruscotti, Age-related changes in cardiac autonomic modulation and heart rate variability in mice, Front Neurosci 15 (2021) 617698. 10.3389/fnins.2021.617698

[37] J.B. Kostis, A.E. Moreyra, M.T. Amendo, J. Di Pietro, N. Cosgrove, P.T. Kuo, The effect of age on heart rate in subjects free of heart disease. Studies by ambulatory electrocardiography and maximal exercise stress test, Circulation 65 (1982) 141–5. 10.1161/01.cir.65.1.141

[38] A.M. Alings, L.N. Bouman, Electrophysiology of the ageing rabbit and cat sinoatrial node--a comparative study, Eur Heart J 14(9) (1993) 1278–88. 10.1093/eurheartj/14.9.1278.

[39] K. Kuga, I. Yamaguchi, Y. Sugishita, I. Ito, Assessment by autonomic blockade of age-related changes of the sinus node function and autonomic regulation in sick sinus syndrome, Am J Cardiol 61(4) (1988) 361–6. 10.1016/0002-9149(88)90945-9

[40] Choi, S., Baudot, M., Vivas, O., & Moreno, C. M. (2022). Slowing down as we age: aging of the cardiac pacemaker’s neural control. GeroScience, 44(1), 1–17. 10.1007/s11357-021-00420-3

[41] S. Rossi, R. Statello, G. Pela, F. Leonardi, A. Cabassi, R. Foresti, G. Rozzi, F.P. Lo Muzio, L. Carnevali, A. Sgoifo, L. Magnani, S. Callegari, P. Pastori, A. Tafuni, D. Corradi, M. Miragoli, E. Macchi, Age-related increases in cardiac excitability, refractoriness and impulse conduction favor arrhythmogenesis in male rats, Pflugers Arch 475 (2023) 731–745. 10.1007/s00424-023-02812-0

[42] J.W. Magnani, N. Wang, K.P. Nelson, S. Connelly, R. Deo, N. Rodondi, E.B. Schelbert, M.E. Garcia, C.L. Phillips, M.G. Shlipak, T.B. Harris, P.T. Ellinor, E.J. Benjamin, A. Health, S. Body Composition, Electrocardiographic PR interval and adverse outcomes in older adults: the health, aging, and body composition study, Circ Arrhythm Electrophysiol 6 (2013) 84–90. 10.1161/CIRCEP.112.975342

[43] S.D. Francis Stuart, L. Wang, W.R. Woodard, G.A. Ng, B.A. Habecker, C.M. Ripplinger, Age-related changes in cardiac electrophysiology and calcium handling in response to sympathetic nerve stimulation, J Physiol 596 (2018) 3977–3991. 10.1113/JP276396

[44] J.R. Stratton, M.A. Pfeifer, J.L. Ritchie, J.B. Halter, Hemodynamic effects of epinephrine: concentration-effect study in humans, J Appl Physiol (1985) 58 (1985) 1199–206. 10.1152/jappl.1985.58.4.1199

[45] C. Jungen, K. Scherschel, C. Eickholt, P. Kuklik, N. Klatt, N. Bork, T. Salzbrunn, F. Alken, S. Angendohr, C. Klene, J. Mester, N. Klocker, M.W. Veldkamp, U. Schumacher, S. Willems, V.O. Nikolaev, C. Meyer, Disruption of cardiac cholinergic neurons enhances susceptibility to ventricular arrhythmias, Nat Commun 8 (2017) 14155. 10.1038/ncomms14155

[46] L. Fabritz, D. Damke, M. Emmerich, S.G. Kaufmann, K. Theis, A. Blana, L. Fortmuller, S. Laakmann, S. Hermann, E. Aleynichenko, J. Steinfurt, D. Volkery, B. Riemann, U. Kirchhefer, M.R. Franz, G. Breithardt, E. Carmeliet, M. Schafers, S.K. Maier, P. Carmeliet, P. Kirchhof, Autonomic modulation and antiarrhythmic therapy in a model of long QT syndrome type 3, Cardiovasc Res 87 (2010) 60–72. 10.1093/cvr/cvq029

[47] K.K. Aras, N.R. Faye, B. Cathey, I.R. Efimov, Critical volume of human myocardium necessary to maintain ventricular fibrillation, Circ Arrhythm Electrophysiol 11 (2018) e006692. 10.1161/CIRCEP.118.006692

[48] A.V. Panfilov, Is heart size a factor in ventricular fibrillation? Or how close are rabbit and human hearts?, Heart Rhythm 3 (2006) 862–4. 10.1016/j.hrthm.2005.12.022

[49] P. Boksa, M. Quik, J.B. Mitchell, B. Collier, W. O’Neil, R. Quirion, Pharmacological activity of N-methyl-carbamylcholine, a novel acetylcholine receptor agonist with selectivity for nicotinic receptors, Eur J Pharmacol 173 (1989) 93–108. 10.1016/0014-2999(89)90012-5

