## Supplementary material for "Age-related structural and functional changes of the intracardiac nervous system": Saasu_E, Supplementary info

### Supplementary Figures

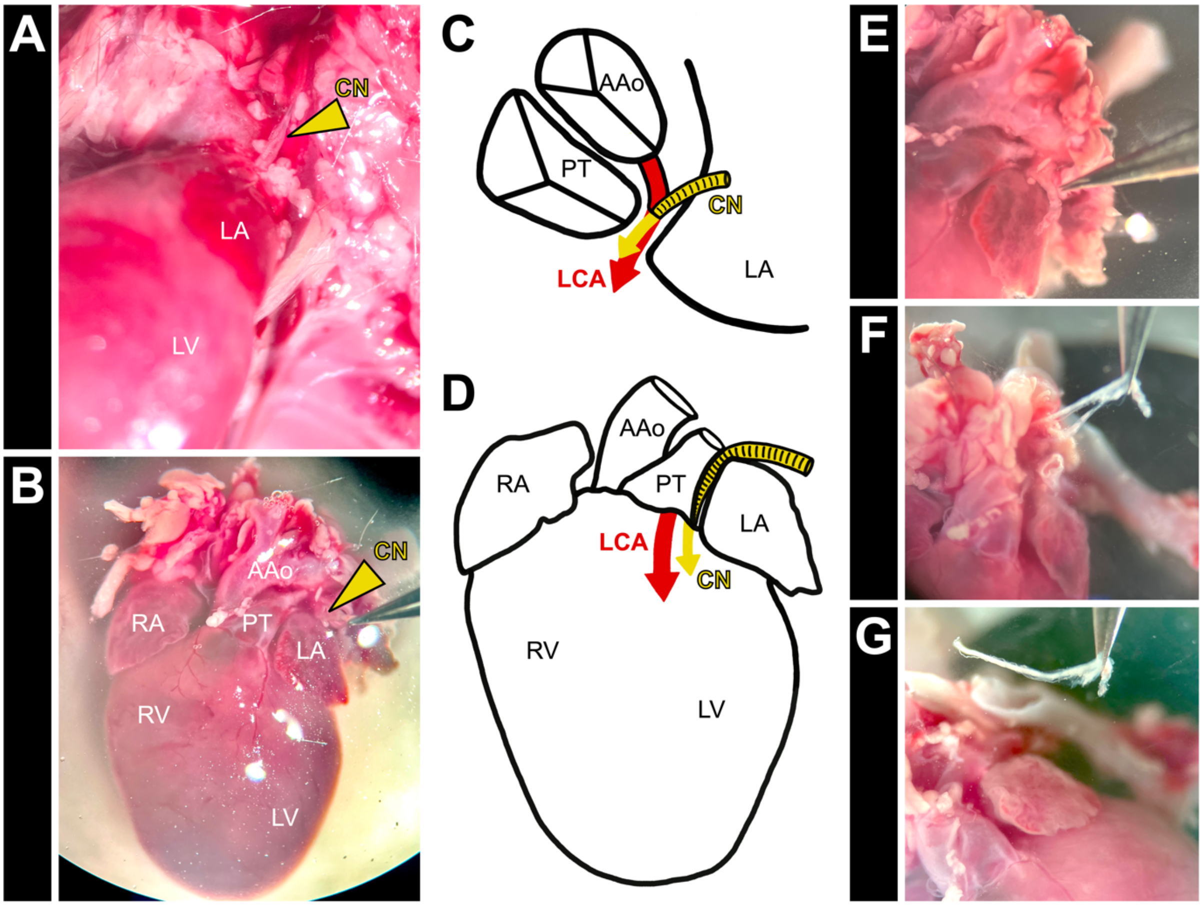


**Figure S1. Schematic overview of cardiac nerve isolation.** (A, B) Path of cardiac nerves (CN, marked with arrowhead) to the heart. Visualization with a dissection microscope in situ (A) and after extraction of the heart (B). After removal of the thymus and lungs, nerves can be displayed running between the ascending aorta (AAo), pulmonary trunk (PT), and the left atrium (LA). (C, D) Schematic representation of the nerves' entrance into the cardiac parenchyma. These fibers appear to enter the ventricles, and run alongside the left coronary artery (LCA). (E, F, G) After visualizing and confirming their path (E), fibers are freed by gently pulling (F) while peeling away surrounding tissue. Long sections of cardiac nerves (G) are isolated. CN: cardiac nerve, LCA: left coronary artery, LA: left atrium, RA: right atrium, AAo: ascending aorta, PT: pulmonary trunk, LV: left ventricle, RV: right ventricle

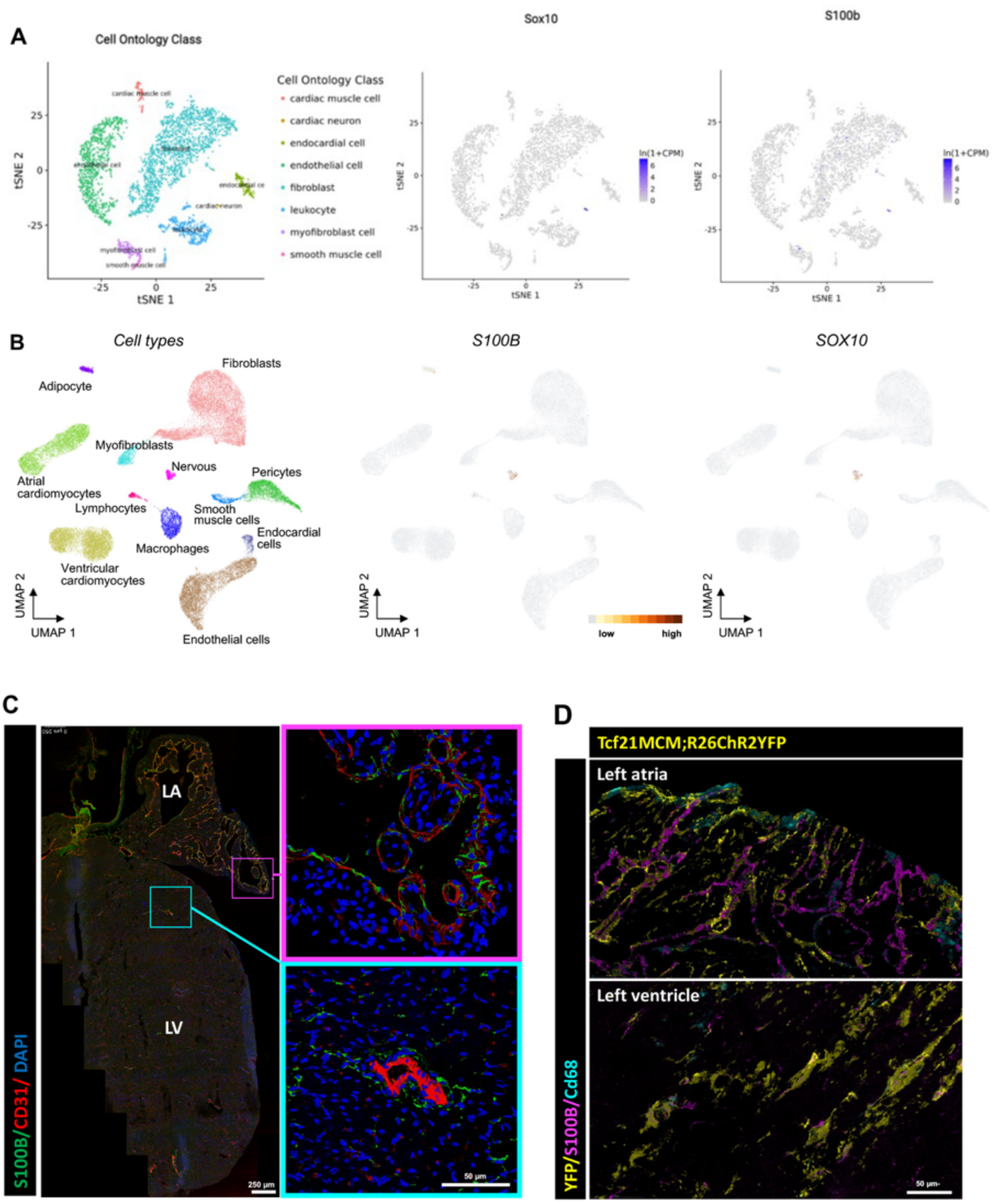


**Figure S2. In the heart, S100B is a specific marker of cardiac SC.** (A) t-SNE representation derived from the online data base *Tabula Muris* shows how in mice, S100B expression is limited to the neural cell-cluster in the heart, matching Sox10 expressing cells. (B) UMAP representation derived from the online data base CARE shows how in human, S100B expression is reduced to the nervous cell-cluster in the heart, matching Sox10 expressing cells. (C) (Left) Representative picture of S100B and the endothelial/endocardial cell marker CD31 expression in a 3-month-old mouse left atria (LA) and left ventricle (LV). (Right) High magnification of atrial endocardium (top, magenta frame) and different sized ventricular blood vessels (bottom, cyan frame). Cardiac SC are localized in intimate contact with endocardial and endothelial cells, matching the areas where nerves can be found in the heart, but there is no co-localization of CD31 and S100B (D) Representative 3D rendering of S100B and the macrophage marker CD68 expression in a 3 month-old *Tcf21*$^{MCM:R26ChR2:YFP}$ mouse left atria (LA) and left ventricle (LV). In this model epicardial cells and cardiac fibroblasts are fluorescently marked with YFP. No co-localization of CD68 or YFP was found in the S100B+ cells, confirming that macrophages, cardiac fibroblasts and epicardial cells, do not express S100B.

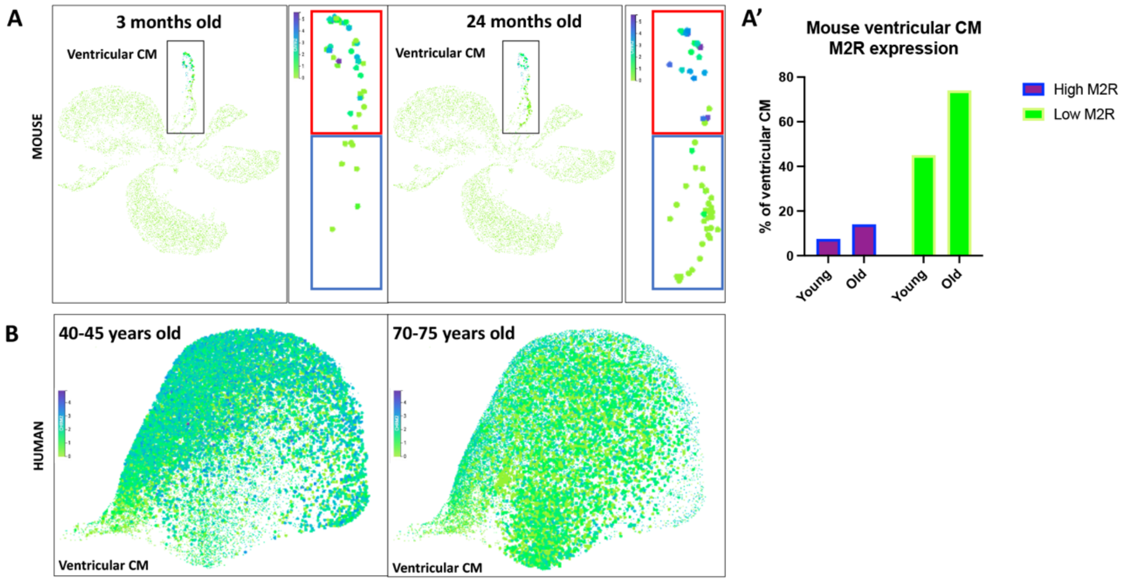


**Figure S3. There are more M2R low-expressing ventricular CM in aged mice and humans.** (A) UMAP representation derived from the online data base *Tabula Muris Senis* shows how in young mice, there are more ventricular CM with high expression of M2R than in young mice. (A') Prevalence of M2R high and low expressing (4-5 and 0-1 from the intensity bar in (A), respectively) ventricular CM in young (3 months old) and old (24 months old) mice. (B) UMAP representation derived from the online data base *Tabula Muris Senis* shows how in 40-45 years old human samples, there are more ventricular CM with high expression of M2R than in 70-75 years old.

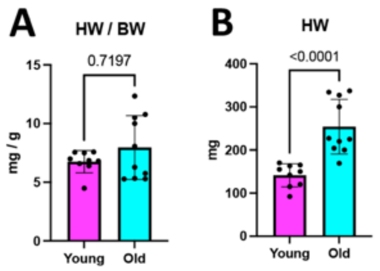


**Figure S4. Hearts from older mice are heavier, but there is no significant difference in the heart weight / body weight ratio.** (A) Heart weight (HW) to body weight (BW) and (B) HW comparisons for old and young mice.

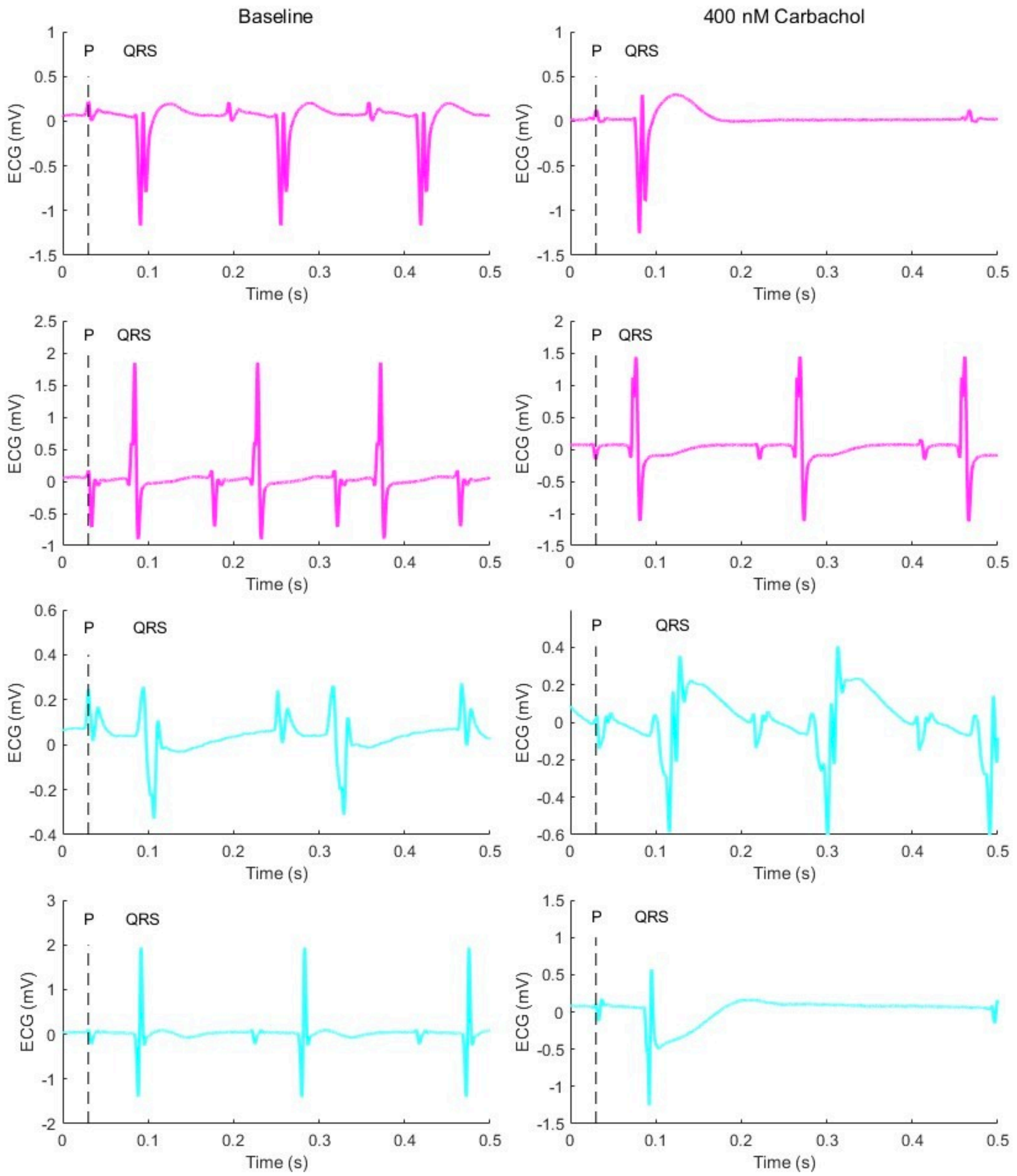


**Figure S5. Representative ECG recordings from Langendorff-perfused hearts from old and young mice.** Recordings in the top two rows are from young (magenta), and in the lower two rows from old mice (cyan). Recordings in the left column show basal conditions, while the those in the right column were taken during expose to 400 nM carbachol. P-wave is always aligned at 0.03 s and shows at least one P-P cycle. P and QRS are labelled for the first cycle.
